# Pulmonary pressure load shapes right ventricular molecular remodelling in dilated cardiomyopathy

**DOI:** 10.64898/2026.08.28.747817

**Authors:** Gino Andrea Bonazza, Aleksandra Paterek, Aleksandra Bogucka, Weronika Ścibek-Rejmontowska, Andrea Laimbacher, Przemysław Leszek, Oliver Distler, Michał Mączewski, Przemysław Błyszczuk, Gabriela Kania

## Abstract

Right ventricular (RV) adaptation to pulmonary hypertension determines outcome in dilated cardiomyopathy (DCM), but the molecular mechanisms of the transition to decompensation remain unclear. We analysed RV tissue from explanted hearts of patients with end-stage DCM using single-nucleus RNA sequencing (n=21), mass spectrometry and Olink Reveal proteomics (both n=44), and integrated these molecular profiles with echocardiographic and right-heart catheterisation measures to identify molecular correlates of RV dysfunction. Mean pulmonary arterial pressure was the dominant correlate of RV transcriptional remodelling, particularly in cardiomyocytes, where higher pressure was associated with contractile remodelling, autophagy, vesicle trafficking and glucose metabolism. In contrast, RV decompensation was characterised by immune activation and reduced oxidative phosphorylation exclusively at the proteomic level. Integrative multi-omics factor analysis (MOFA) further identified fibrosis as the dominant molecular program shared across transcriptomic and proteomic layers. Together, these findings indicate molecular adaptation to pressure load and tissue fibrosis during progression towards RV failure.

## Introduction

Right ventricular (RV) function is a major determinant of clinical outcome in pulmonary hypertension (PH). Prognosis in PH is closely linked to the capacity of the RV to preserve pump function despite increasing pulmonary afterload, yet the mechanisms that determine whether the RV adapts or progresses toward failure remain incompletely understood^1–4^.

PH due to left heart disease, including advanced dilated cardiomyopathy (DCM), develops when elevated left ventricular filling pressures are transmitted backwards to the pulmonary circulation^5–7^. This initially causes post-capillary PH, but sustained elevation of pulmonary pressure may be accompanied by pulmonary vasoconstriction and vascular remodelling, increasing pulmonary vascular resistance (PVR) and imposing a chronic pressure load on the RV^5,6^. As mean and systolic pulmonary arterial pressures (mPAP and sPAP) and PVR increase, the RV may preserve pump function through tissue hypertrophy leading to enhanced contractility, and structural remodelling^1–3,8^. This compensatory response can preserve RV systolic function to increased pulmonary pressure for some time. This mechanism is known as RV–pulmonary arterial (RV–PA) coupling and reflects the ability of the right ventricle to adapt its contractility to the afterload imposed by the pulmonary circulation^3,4,8^. Although the gold-standard assessment of RV–PA coupling is the ratio of end-systolic ventricular elastance to arterial elastance (Ees/Ea), it can be estimated non-invasively using the tricuspid annular plane systolic excursion (TAPSE)/systolic pulmonary arterial pressure (sPAP) ratio^4,9,10^. With persistent or excessive increases in afterload, however, compensatory adaptation may become insufficient. Progressive RV hypertrophy and remodelling are then accompanied by impaired contractile reserve, RV–PA uncoupling, and eventual RV dilation and systolic dysfunction. As a consequence, right-sided filling pressures rise, systemic venous congestion develops, and clinical outcomes worsen^1–3,8^.

The pathophysiology of RV failure differs from that of left ventricular failure. The RV has distinct anatomical, metabolic and biomechanical properties and is particularly sensitive to changes in afterload. Accordingly, therapeutic strategies that improve outcomes in left ventricular systolic failure have not shown comparable benefit in RV failure associated with PH, highlighting the need for molecular studies focused directly on the human RV^1–3,8^.

Omics approaches have substantially advanced the understanding of left ventricular remodelling in heart failure, but the human RV remains far less characterised^11–13^. Existing molecular studies of RV remodelling have often relied on experimental models or focused on selected pulmonary vascular diseases, while PH due to left ventricular disease remains comparatively understudied despite being one of the most common forms of PH. Moreover, experimental studies allow RV tissue to be sampled before and after the development of pressure overload and across stages of adaptation and decompensation. In humans, however, RV tissue is usually available only at surgery, transplantation or autopsy, limiting direct temporal reconstruction of this process^14–17^.

In this study, we analysed human RV free-wall samples from patients with advanced DCM, a clinically common setting of PH due to left ventricular disease. By integrating single-nucleus RNA sequencing and tissue proteomics with invasive haemodynamic and echocardiographic measurements, we aimed to define molecular changes associated with the continuous process from adaptation to pressure overload to RV dysfunction and RV-pulmonary arterial uncoupling.

## Results

### Transcriptional profiling of right ventricular tissue from DCM patients

To investigate how the RV adapts and becomes dysfunctional in PH associated with left heart disease, we performed snRNA-seq on snap-frozen specimens from the RV free wall (Fig. 1A-B) of 21 male patients with DCM. All patients had end-stage heart failure and underwent heart transplantation. As expected, patients exhibited severely impaired left ventricular ejection fraction (LVEF; mean 16.8% ± 6.7) and markedly enlarged LV (mean end-diastolic diameter 77.5 mm ± 11.7). Notably, 11 of the 21 DCM patients had reduced RV contractility, as indicated by a tricuspid annular plane systolic excursion (TAPSE) < 16 mm. Right ventricular dimensions were also frequently increased, with 12 of 21 patients showing an RV inflow tract dimension (RVIT) > 42 mm. Additionally, 17 of 18 patients who underwent RV catheterisation met the haemodynamic criteria for PH (mPAP > 20 mmHg)^5^, with 16 patients showing also PCWP > 15 mmHg and 14 patients showing PVR > 2 Wood Units (WU) (Fig. 1C, Supplementary Data 1). Correlation analysis showed strong correlations among PH-related haemodynamic parameters, including mPAP, sPAP, PCWP, and PVR, and inverse relationships between RV or LV functional and structural measurements (Extended Data Fig. 3A).

**Fig. 1:**
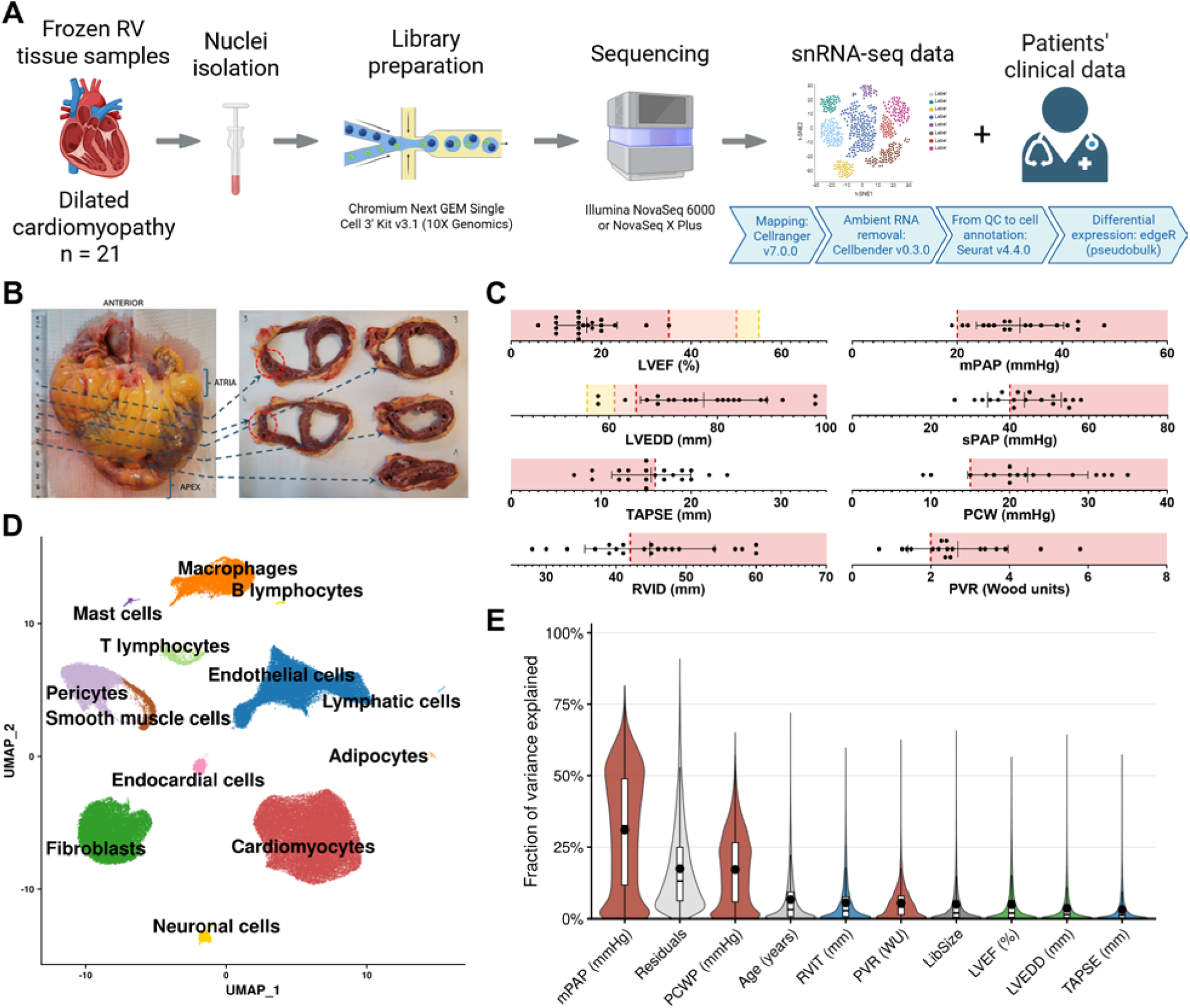
Clinical characteristics of the patient cohort and single-nuclei transcriptomic profiling of the right ventricle. **A,** Overview of the study design. **B,** Preparation of tissue samples from explanted hearts. Red circles indicate regions of the RV harvested for analysis. **C,** Selected echocardiographic (LVEF, LVEDD, RVIT, TAPSE) and haemodynamic (mPAP, sPAP, PCWP, PVR) parameters of the DCM cohort. Abnormal ranges are highlighted in red (n = 21). **D,** Uniform manifold approximation and projection (UMAP) of the thirteen cell types identified in the dataset. **E,** Variance partitioning analysis performed on sample level pseudobulk profiles from all nuclei. mPAP explained the largest fraction of gene expression variance among the clinical variables tested, followed by PCWP. A large fraction of variance remained unexplained by the clinical variables included in the model, as reflected by the residual component.

Following quality control, a total of 136,769 nuclei were retained for downstream analysis. Dimensionality reduction and unsupervised clustering identified 13 distinct cell types, each defined by canonical marker genes (Extended Data Fig. 1, Extended Data Fig. 2, Supplementary Data 2). These included cardiomyocytes, fibroblasts, endothelial cells, pericytes, macrophages, T lymphocytes, smooth muscle cells, neuronal cells, endocardial cells, adipocytes, mast cells, lymphatic cells and B lymphocytes (Fig. 1D).

### Transcriptional changes in the right ventricle of DCM patients are primarily associated with the severity of pulmonary hypertension

To identify which clinical variables account for the largest share of variance in gene expression, we performed variance partitioning on pseudobulk profiles pooling all cell types (Fig. 1E). The model included TAPSE, RVIT, LVEF, LVEDD, mPAP, PCWP, PVR, age and library size. mPAP accounted for the largest fraction of explained variance, followed by PCWP. Age and library size, two potential confounders, contributed modestly. As expected, a substantial residual variance remained. These results indicate that PAP is the dominant clinical correlate of RV gene expression variation in this cohort.

We next tested linear relationships between gene expression levels and clinical variables of interest, including PH-related haemodynamic parameters (mPAP, sPAP, PCWP, and PVR), RV functional and structural parameters (TAPSE and RVIT), and LV functional and structural parameters (LVEF and LVEDD). For each cell type, we performed pseudobulk differential expression analysis using quasi-likelihood negative binomial generalized linear models, fitting separate models for each parameter and including age as a scaled covariate. All tested variables were modelled as continuous predictors and scaled, allowing direct comparison of log₂ fold changes (log₂FC) across parameters. The resulting numbers of positively and negatively associated genes for each parameter-cell type combination are summarized in separate heatmaps (Fig. 2A-B). The strongest associations were observed for mPAP. This was most pronounced in cardiomyocytes, where 970 genes were positively and 2,218 genes negatively associated with mPAP. Other cell types showed fewer transcriptional changes, with mPAP-associated genes also observed in fibroblasts, endothelial cells and pericytes. sPAP and PVR showed similar but weaker patterns, whereas PCWP showed no significant associations. In contrast, metrics of RV function or size, including TAPSE and RVIT, showed almost no significant gene expression changes in any major cell type. LV parameters showed only minor associations, mainly with LVEDD. We also evaluated cell type abundance using differential abundance models, but no major changes in the proportion of any main cell population were associated with the tested parameters (Supplementary Data 4).

**Fig. 2:**
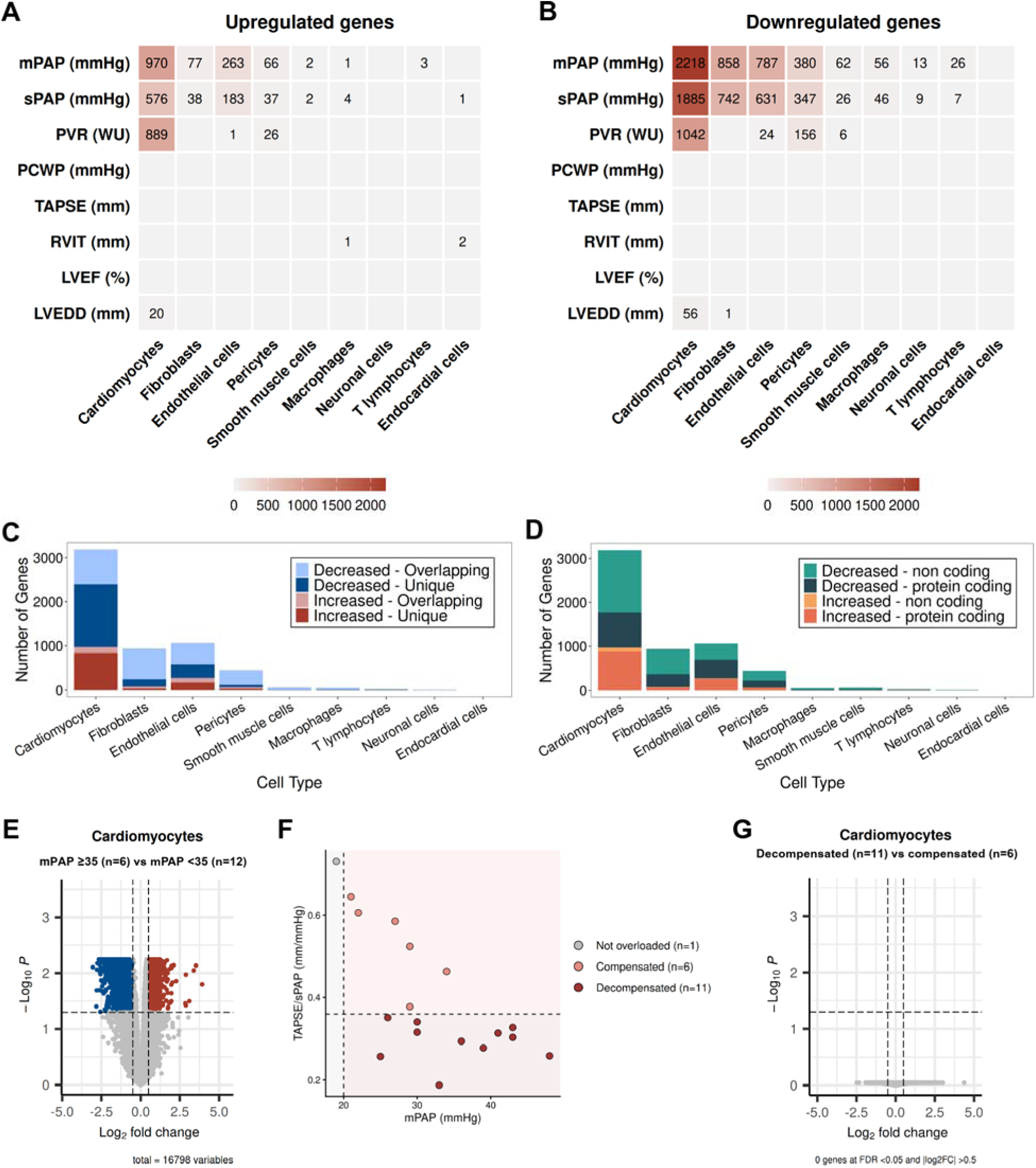
Cell type-resolved differential expression analysis using echocardiographic and haemodynamic parameters in RV tissue. **A-B**, Heatmaps showing the number of genes positively (**A**) and negatively (**B**) associated with each clinical parameter in each major cell type. Differential expression was dominated by PH-related haemodynamic parameters, particularly mPAP, with the largest number of associated genes detected in cardiomyocytes. sPAP and PVR showed similar but weaker patterns, whereas RV functional and structural parameters showed almost no significant associations. **C**, Stacked bar plots showing, for each cell type, the number of genes significantly associated with mPAP (FDR < 0.05, |log₂FC| > 0.5). Bars are partitioned into genes unique to a given cell type (dark red and dark blue for increased and decreased expression, respectively) and genes shared by at least two cell types (lighter red and blue). **D**, Stacked bar plots depicting, for the same set of mPAP-associated genes, the contribution of different gene biotypes in each cell type. Protein-coding genes are shown in dark shades, whereas non-coding RNAs are represented by the corresponding lighter tones. **E**, Volcano plot showing cardiomyocyte differential expression between patients with mPAP ≥35 mmHg (n = 6) and mPAP <35 mmHg (n = 12). Positive log₂FC values indicate higher expression in the mPAP ≥35 mmHg group. **F**, Classification of patients according to pulmonary pressure load and RV–pulmonary arterial coupling. Points represent patients with available mPAP and TAPSE/sPAP measurements. Patients with mPAP ≤20 mmHg were classified as not pressure-overloaded (n = 1). Patients with mPAP >20 mmHg and TAPSE/sPAP >0.36 were classified as compensated (n = 6), whereas those with mPAP >20 mmHg and TAPSE/sPAP ≤0.36 were classified as decompensated (n = 11). Dashed lines indicate the thresholds used for classification. **G**, Volcano plot showing cardiomyocyte differential expression between decompensated (n = 11) and compensated (n = 6) patients. Positive log₂FC values indicate higher expression in the decompensated group.

Since mPAP showed the strongest transcriptional associations across cell types, we further characterised the mPAP-associated genes from the continuous models. Interestingly, a substantial fraction of these genes was shared across multiple cell types (Fig. 2C), potentially reflecting a common transcriptional response. Additionally, non-coding transcripts accounted for 10.1% of upregulated genes and 64.7% of downregulated genes in cardiomyocytes, highlighting a potential regulatory role of non-coding RNAs in RV remodelling under increased pressure load (Fig. 2D).

To test whether the same patterns were observed when patients were divided into clinically interpretable groups, we repeated the analysis using predefined thresholds. This analysis was performed in cardiomyocytes, where the mPAP-associated transcriptional signal was strongest. For mPAP, almost all patients were above the current diagnostic threshold for PH (mPAP >20 mmHg)^5^, making this cutoff uninformative for subgroup analysis. We therefore compared patients with markedly elevated mPAP (≥35 mmHg) to those with lower mPAP (<35 mmHg) (Fig. 2E). This comparison gave results that were very similar to the continuous mPAP model (Extended Data Fig. 3B-C). In total, 72.6% of genes associated with mPAP in the continuous model were also significant in the mPAP ≥35 mmHg comparison, with 554 shared upregulated genes and 1,761 shared downregulated genes. The log₂FC estimates were strongly correlated between the two approaches (Pearson r = 0.97), indicating that the threshold-based analysis captured the same mPAP-associated transcriptional program.

We next classified patients according to RV–pulmonary arterial coupling. Among patients with mPAP >20 mmHg, those with TAPSE/sPAP >0.36 were classified as compensated and those with TAPSE/sPAP ≤0.36 as decompensated (Fig. 2F). One patient with mPAP ≤20 mmHg was classified as not pressure-overloaded and excluded from this comparison. No significant differentially expressed genes were identified between decompensated and compensated patients in cardiomyocytes (Fig. 2G). These findings further indicate that RV cardiomyocyte gene expression in this cohort primarily reflects pulmonary pressure load rather than a distinct transcriptional state associated with RV decompensation.

### Cardiomyocytes exhibit profound transcriptional changes in association to elevated pulmonary arterial pressure

To identify dysregulated pathways in cardiomyocytes, we performed gene set enrichment analysis using Gene Ontology (GO) terms (Fig. 3A) and KEGG pathways (Fig. 3B). Analysis of GO terms and pathways networks (Fig. 3C-D) revealed that cardiomyocytes adapt to elevated PAP by upregulating gene programs involved in sarcomere remodelling, autophagy, endosomal transport, adrenergic signalling, and glucose metabolism. Gene set scores derived from the aggregated expression of genes in these processes showed strong positive correlations with mPAP, supporting their functional relevance (Fig. 3F).

**Fig. 3:**
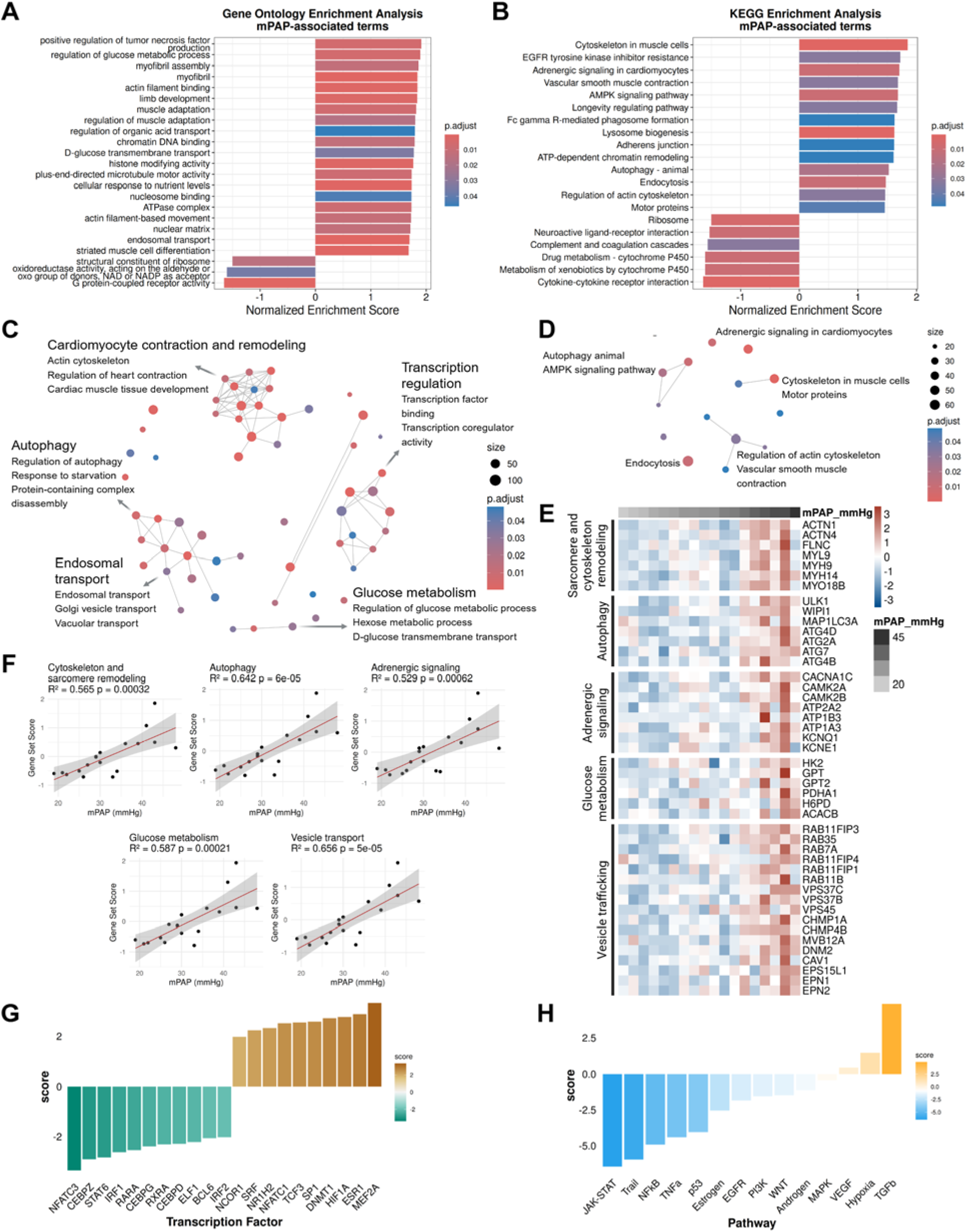
Gene programs associated with elevated pulmonary arterial pressure in RV cardiomyocytes. **A,** Bar plot showing the top 20 Gene Ontology (GO) terms enriched among mPAP-associated genes in cardiomyocytes, ranked by the absolute value of the normalized enrichment score (NES). **B,** Equivalent bar plot for KEGG pathways, displaying the 20 pathways with the highest absolute NES among mPAP-associated genes. **C,** Network visualisation of enriched GO terms among mPAP-associated genes. Each node represents a GO term, edges connect pairs of terms sharing more than 20% of their genes. Clusters of densely connected terms were manually annotated with summary labels (“Transcription regulation”, “Cardiomyocyte contraction and remodelling”, “Autophagy”, “Endosomal transport”, “Glucose metabolism”), and a few representative GO terms are shown for each cluster. **D,** Network visualisation of enriched KEGG pathways, with nodes and edges defined as in panel C. **E**, Heatmap of representative mPAP-associated genes grouped according to the main functional programs identified. Columns correspond to individual DCM samples ordered by increasing mPAP, and rows represent genes; colours indicate scaled expression values. **F,** Scatter plots showing the relationship between mPAP and gene-set scores for the key functional processes identified. A gene-set score summarising the expression of all genes in the set was computed per sample and plotted against mPAP. Lines represent linear regression fits with 95% confidence intervals; R² and P values for the regression are reported in each panel. **G**, Transcription factor activity inference showing top candidate regulators positively (brown) and negatively (green) associated with mPAP-related gene expression changes. **H,** Pathway activity inference indicating signalling cascades predicted to be activated (orange) or suppressed (cyan) in association to increased pulmonary pressure.

Among the genes with increased expression at higher mPAP, we identified key components of the cytoskeletal and sarcomeric structure (*ACTN1, ACTN4, FLNC, MYL9, MYH9, MYH14, MYO18B*) suggesting a profound remodelling of the contractile machinery in cardiomyocytes (Fig. 3E). The upregulation of autophagy-related genes included the core autophagy initiation kinase *ULK1*, genes involved in autophagosome formation (*WIPI1, MAP1LC3A*), and additional autophagy regulators (*ATG4D, ATG2A, ATG7, ATG4B*). Among genes involved in calcium handling and ion transport, we observed significant upregulation of CACNA1C, CAMK2A, CAMK2B and ATP2A2, together with other ion transporters including ATP1B3, ATP1A3, KCNQ1 and KCNE1. Increased expression of genes in the glycolytic pathway (*HK2, GPT, GPT2, PDHA1, H6PD*), along with the negative regulator of fatty acid oxidation *ACACB*, suggests a metabolic shift favouring glucose metabolism. Additionally, many genes positively associated with mPAP were involved in vesicle trafficking (*RAB11FIP3, RAB35, RAB7A, RAB11FIP4, RAB11FIP1, RAB11B*), including members of the endosomal complex required for transport (*VPS37C, VPS37B, VPS45, CHMP1A, CHMP4B, MVB12A*) and genes associated with endocytosis (*DNM2, CAV1, EPS15L1, EPN1, EPN2*), highlighting alterations in intracellular trafficking and vesicle dynamics in response to increasing PAP.

To address the molecular mechanisms driving these transcriptional changes, we used computational tools to infer transcription factor and pathway activity. Among the most active transcription factors identified were myocyte enhancer factor 2A (MEF2A) and serum response factor (SRF), both of which play critical roles in cardiac development and pathology (Fig. 3G). Hypoxia-inducible factor-1α (HIF-1α), another top transcription factor, plays a key role in pressure overload-induced cardiac remodelling, supporting cardiomyocyte function by preserving calcium handling and contractility. Pathway analysis revealed significant activation of hypoxia and TGF-β signalling in cardiomyocytes, the latter being a key driver of interstitial fibrosis in response to pressure overload, promoting extracellular matrix deposition and stiffening of the myocardium (Fig. 3H).

### Non-myocyte transcriptional programs associated with pulmonary pressure load

To characterize transcriptional programs associated with pulmonary hypertension severity across non-myocyte RV cell types, we performed GSEA on ranked mPAP-associated pseudobulk differential expression results (Fig. 4A). Endothelial cells showed positive enrichment of small GTPase signalling, together with negative enrichment of respiratory-chain and ribosomal translation terms. Pericytes showed positive enrichment of chromatin-regulatory processes, including SWI/SNF complex, histone modification and chromatin remodelling. Macrophages showed positive enrichment of lysosomal processes and negative enrichment of ribosomal translation. In T lymphocytes, positively enriched terms included lymphocyte activation and antigen-receptor signalling, whereas membrane and exocytosis processes were negatively enriched. Full enrichment results are provided in Supplementary Data 5.

**Fig. 4:**
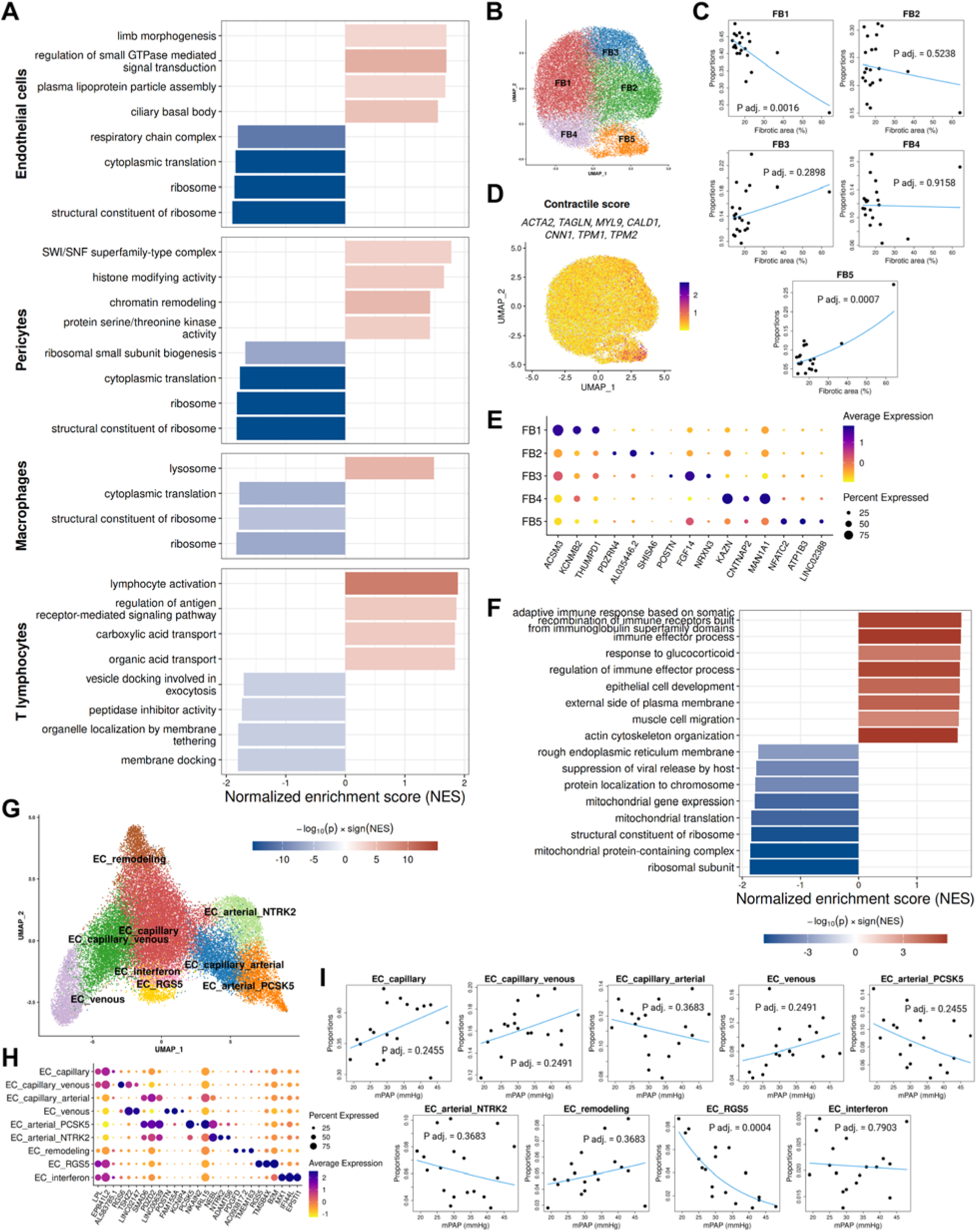
mPAP-associated pathway changes across non-myocyte RV cell types and subset analyses. **A**, Gene set enrichment analysis (GSEA) performed on ranked, mPAP-associated differential expression results (age-adjusted) for major non-myocyte RV cell types. Bar plots show normalized enrichment scores (NES) for selected Gene Ontology terms; colour indicates signed significance (−log10(P) × sign(NES)). **B**, UMAP of RV fibroblasts showing five transcriptionally distinct fibroblast subsets (FB1-FB5). **C**, Association between fibroblast subset proportions and histological fibrosis (fibrotic area, %) across samples. Differential abundance was tested using the propeller method (logit-transformed per-sample proportions) with a limma empirical Bayes linear model, including age as a covariate. **D**, Fibroblast contractile score mapped onto the fibroblast UMAP, derived from the average expression of canonical contractile markers (ACTA2, TAGLN, MYL9, CALD1, CNN1, TPM1, TPM2). **E**, Dot plot summarising fibroblast subset markers, showing the top three marker genes (by logFC) for each subset. Dot size indicates the fraction of nuclei expressing each gene and colour indicates average scaled expression. **F**, GSEA performed on ranked, fibrosis-associated differential expression results in fibroblasts (fibrosis as tested variable, age-adjusted). Bar plots show NES for selected terms; colour indicates signed significance as in (A). **G**, UMAP of RV endothelial cells annotated into nine subsets: EC capillary, EC remodelling, EC capillary-venous, EC venous, EC capillary-arterial, EC arterial-PCSK5, EC arterial-NTRK2, EC-interferon, and EC capillary-RGS5. **H**, Dot plot summarising endothelial subset markers, showing the top three marker genes (by logFC) for each subset. **I**, Association between endothelial subset proportions and mPAP across samples. Differential abundance was tested using the propeller method (logit-transformed per-sample proportions) with a limma empirical Bayes linear model, including age as a covariate.

We next separated major cell types into transcriptionally distinct subsets and tested whether subset proportions were associated with mPAP. Fibroblasts separated into five subsets (Fig. 4B, Extended Data Fig. 5A-B), including a *FAP*- and *POSTN*-high activated population (FB3) and a myofibroblast subset (FB5) marked by genes such as *PDLIM3* and *GPC6* (Fig. 4E). FB5 also showed higher expression of contractile genes, captured by a contractile score based on *ACTA2*, *TAGLN*, *MYL9*, *CNN1*, *CALD1*, *TPM1*, and *TPM2* (Fig. 4D). No fibroblast subset proportion was significantly associated with mPAP (Extended Data Fig. 8A, Supplementary Data 6).

Cardiac fibrosis is enhanced in DCM hearts. Although histological fibrosis measured by Masson’s trichrome in RV tissues did not correlate with mPAP (Extended Data Fig. 8B), it was positively associated with the FB5 myofibroblast subset and negatively associated with the dominant subset FB1 (Fig. 4C).

Consistent with the contractile phenotype of FB5, GSEA of fibrosis-associated transcriptional changes in fibroblasts showed enrichment of terms related to actin cytoskeleton organisation and muscle cell migration, together with immune-related pathways (Fig. 4F).

Endothelial cells were separated into nine subsets including capillary, arterial, and venous cells (Fig. 4G-H, Extended Data Fig. 5E-F). Only the *RGS5*-positive capillary subset showed a significant negative association with mPAP (Fig. 4I). This subset was characterised by high *RGS5* expression together with capillary endothelial markers such as *FABP4, FABP5, and AQP1*. It also expressed genes related to interferon signalling and antigen presentation, including *B2M, HLA-B, HLA-C, IFI27*, and *IFITM3*, suggesting an immune-activated cell state (Extended Data Fig. 5E-H). Furthermore, subset analyses in other cell types, including cardiomyocytes, macrophages, mural cells, and T lymphocytes, did not show additional mPAP-associated changes in subset abundance and are shown in Extended Data Fig. 4, 6 and 7. Complete marker lists and differential abundance results for the subclustered cell populations are provided in Supplementary Data 3.

**Fig. 7:**
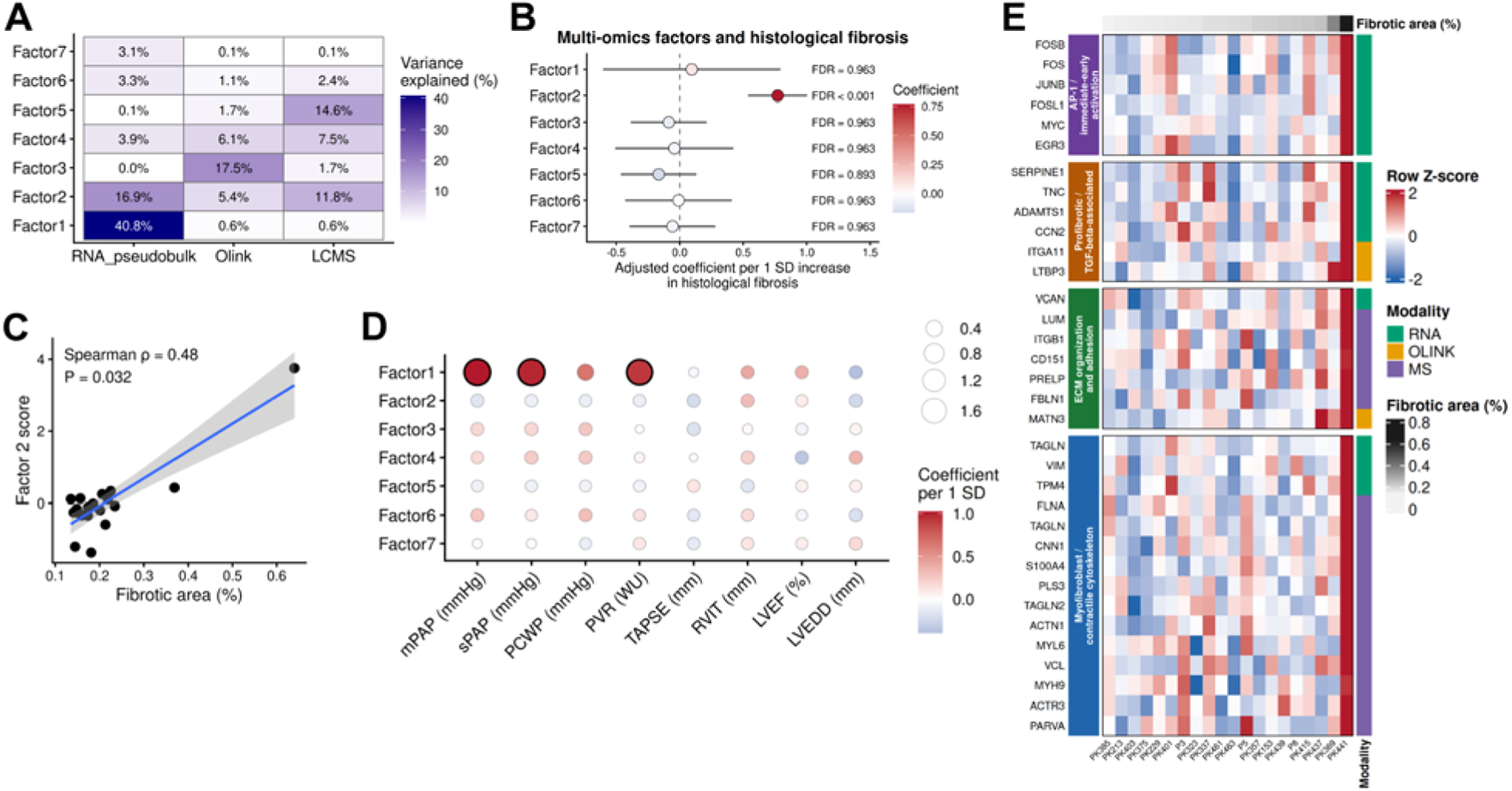
MOFA identifies a fibrosis-associated molecular factor across transcriptomic and proteomic datasets. **A**, Variance explained by the seven factors from the MOFA model integrating sample-level pseudobulk snRNA-seq, Olink and LC-MS/MS data from the 20 patients measured with all three approaches. The RNA pseudobulk dataset consisted of the 2,000 genes with the highest variance after variance-stabilizing transformation of sample-level pseudobulk counts. **B**, Associations between the seven multi-omics factor scores and histological RV fibrosis, measured as the percentage of fibrotic area in Masson’s trichrome-stained sections. Separate linear models were fitted for each factor, with the factor score as the outcome and scaled fibrotic area as the predictor, adjusting for scaled age. Points indicate adjusted coefficients per 1 SD increase in fibrotic area, and horizontal lines show 95% confidence intervals. **C**, Relationship between Factor 2 scores and histological RV fibrosis. Points represent individual patients, and the line shows a linear regression fit with its 95% confidence interval. The Spearman correlation coefficient and corresponding P value are reported. **D**, Associations between the multi-omics factor scores and continuous haemodynamic and echocardiographic parameters. Separate linear models were fitted for each factor-parameter combination, with adjustment for scaled age. Colour indicates the adjusted coefficient per 1 SD increase in the clinical parameter, and dot size represents −log₁₀(FDR). Black outlines indicate associations with FDR < 0.10. **E**, Heatmap of representative features selected among the top 100 features with the highest Factor 2 weights across the RNA, Olink and LC-MS/MS datasets and annotated according to their biological functions. Rows show gene expression or protein abundance standardised as Z-scores across samples. Columns represent patients ordered by increasing fibrotic area. The top annotation shows histological fibrotic area, and the side annotation indicates the molecular modality of each feature.

### RV pressure overload is associated with TGF-β-mediated cell-cell interactions

To investigate how intercellular communication changes with PH severity, we stratified patients into mPAP≥35 and mPAP<35 groups and computed differential interactions using MultiNicheNet. Overall, most top-ranked interactions were enriched in the mPAP≥35 group (Fig. 5A). Prioritized interactions involved multiple cell types, with frequent contributions from endothelial cells, fibroblasts and T lymphocytes. The MultiNicheNet prioritization network (Fig. 5B) highlighted a fibroblast-centred hub of TGF-β signalling in the mPAP≥35 group, where interactions between TGFB1/TGFB3 and TGF-β receptors converge on SERPINE1, a canonical effector of profibrotic remodelling. This core axis was connected to endothelial-immune and vascular interactions, including CSF1–CSF1R, ITGAL–ICAM1 and GLG1–UNC5B. Other relevant interactions enriched in the mPAP≥35 group included SERPINE1–ITGAV and THBS1–SDC4 from fibroblasts to cardiomyocytes, and JAG2–NOTCH3 from endothelial cells to pericytes (Fig. 5C-D). Together, these results point to a dense fibroblast-centred network in samples with markedly high mPAP that links profibrotic TGF-β activity with adhesion and immune-vascular interactions, driving maladaptive RV remodelling under pressure overload.

**Fig. 5:**
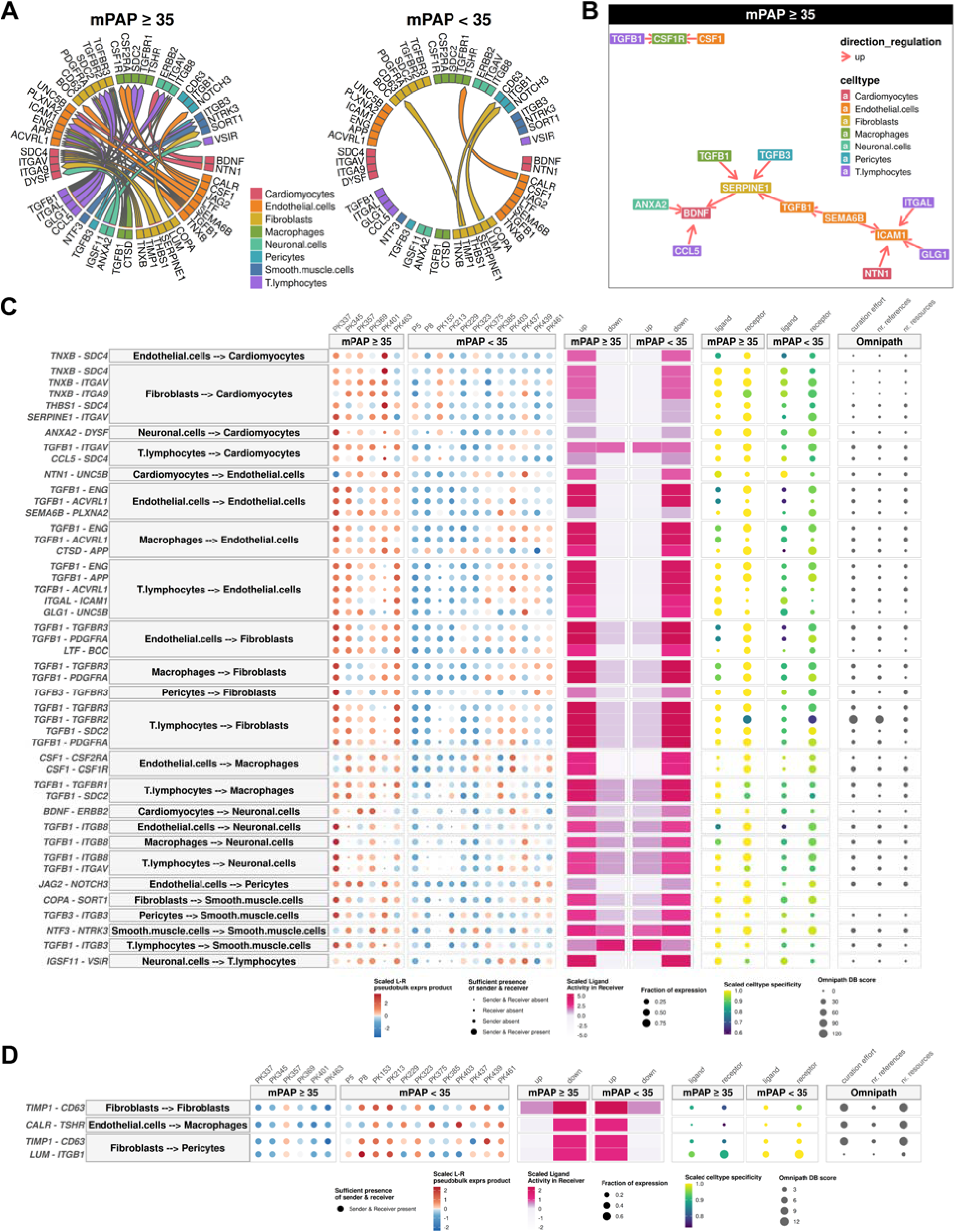
mPAP-related changes in intercellular communication inferred by MultiNicheNet. **A**, Chord diagrams showing the top prioritised ligand-receptor (LR) interactions in mPAP≥35 and mPAP<35 groups. Arcs connect ligands (bottom) to their corresponding receptors (top), with arrowheads indicating the direction of signalling from sender to receiver cell type and colours denoting the sender cell type. **B**, MultiNicheNet intercellular regulatory network for the mPAP≥35 group, highlighting a fibroblast-centred TGF-β signalling module. Nodes represent ligands, receptors and the downstream target gene SERPINE1, and are coloured according to the predominant expressing cell type. Directed edges depict putative signalling links from ligands in sender cells to receptors in receiver cells and further to the target gene; arrowheads indicate the inferred direction of regulation. **C-D**, Bubble heatmaps summarising the highest-ranked LR pairs in mPAP≥35 (**C**) and mPAP<35 (**D**) conditions. Rows correspond to specific sender-receiver LR combinations. The left panels show scaled ligand-receptor pseudobulk expression products across individual samples in mPAP≥35 and mPAP<35 groups. The right panels summarise the evidence features used for MultiNicheNet prioritisation: sufficient presence of sender and receiver cell types, scaled ligand activity in the receiver cell type, fraction of expressing cells, cell-type specificity and Omnipath database support.

### The RV tissue proteome is associated with RV function rather than pulmonary pressure load

To test whether the transcriptional programs associated with pulmonary pressure load were also reflected at the protein level, we performed proteomic profiling of the full 44-patient DCM cohort, which included all 21 patients analysed by snRNA-seq. The correlation structure of clinical variables in the extended proteomics cohort closely mirrored that observed in the snRNA-seq cohort (Extended Data Fig. 9A). We analysed the samples using two complementary approaches, liquid chromatography tandem mass spectrometry (LC-MS/MS) and the Olink Reveal panel. LC-MS/MS detected 705 proteins and mainly covered abundant tissue proteins, whereas the Olink Reveal panel included 1,034 proteins and covered several lower-abundance and secreted proteins (Fig. 6A). The two platforms exhibited largely complementary coverage, with only 41 proteins detected in both datasets. After exclusion of low-quality samples, the number of proteins detected by LC-MS/MS was similar across analysed samples, ranging from 556 to 671 proteins per sample (Extended Data Fig. 9B). Variance partitioning showed that the clinical variables explained only a small proportion of the overall variation in either proteomic dataset (Extended Data Fig. 9C-D).

**Fig. 6:**
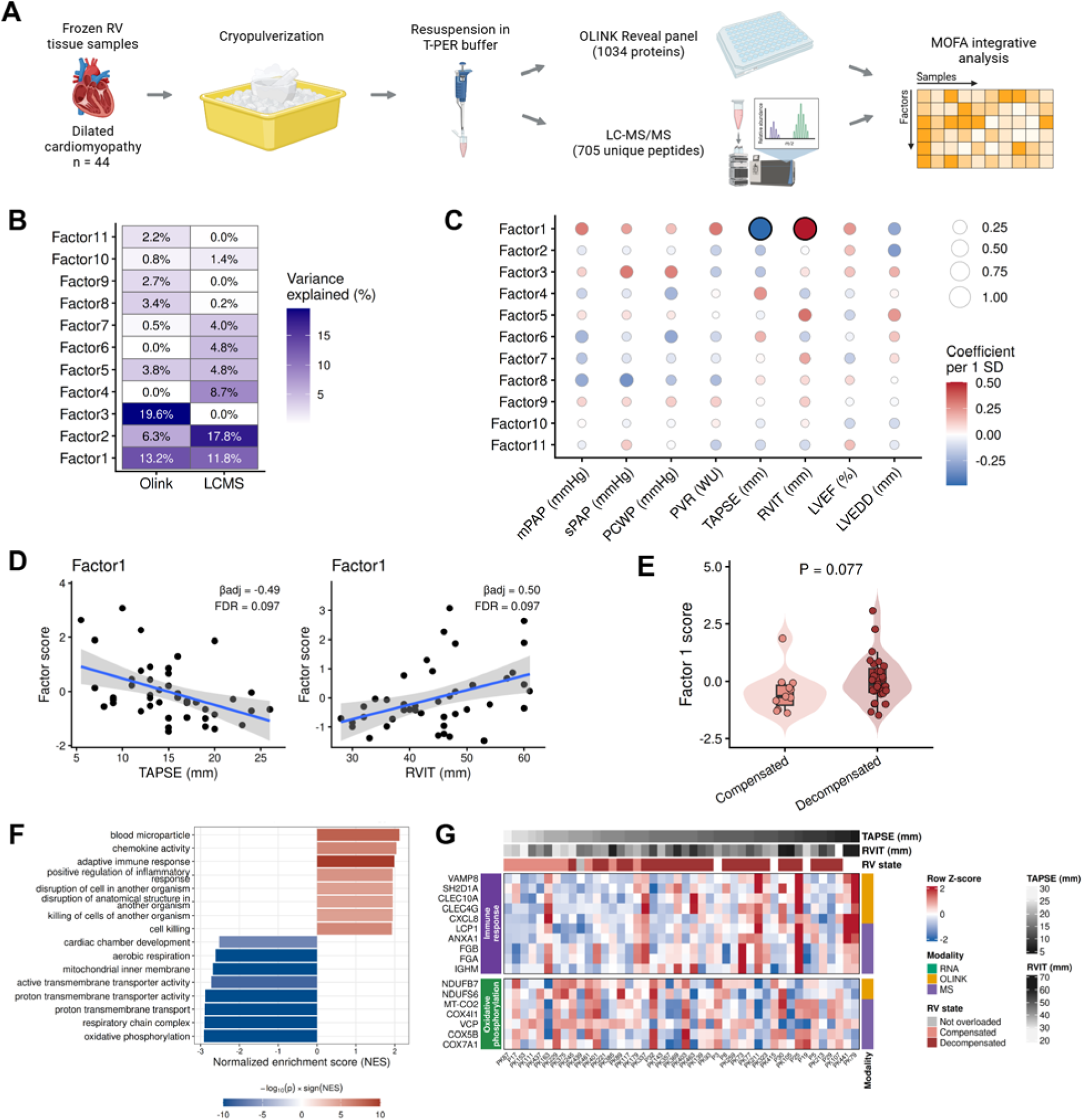
Proteomic variation is associated with RV dysfunction and decompensation. **A**, Overview of the proteomic workflow. Frozen RV free-wall tissue from 44 patients with DCM was cryopulverized and lysed in T-PER buffer. Protein abundance was measured using the Olink Reveal panel and LC-MS/MS, and the two datasets were jointly analysed using Multi-Omics Factor Analysis (MOFA). The Olink dataset contained 1,034 proteins, whereas the LC-MS/MS dataset contained 705 proteins before feature filtering. For MOFA, Olink values were adjusted for differences in sample protein concentration, and LC-MS/MS proteins detected in at least 50% of samples were retained. **B**, Heatmap showing the percentage of variance in each proteomic dataset explained by the 11 MOFA factors. The number of factors was selected by fitting models with 5-15 factors using three random seeds. For each number of factors, the model with the highest evidence lower bound was retained. The final factor number was selected when the weakest additional factor explained less than 2% of the combined variance across views in three consecutive models. **C**, Associations between MOFA factor scores and continuous haemodynamic and echocardiographic parameters. Separate linear models were fitted for each factor– parameter combination, with the factor score as the outcome and the scaled clinical parameter as the predictor, adjusting for scaled age. Colour indicates the adjusted coefficient per 1 SD increase in the clinical parameter, and dot size represents −log₁₀(FDR). Black outlines indicate associations with FDR < 0.10. **D**, Relationships between Factor 1 scores and TAPSE and RVIT. Points represent individual patients, and lines show unadjusted linear regression fits with 95% confidence intervals. **E**, Factor 1 scores in patients with compensated and decompensated RV function. Factor scores were compared using a linear model adjusted for age. **F**, Gene Ontology gene set enrichment analysis of proteins ranked by their Factor 1 weights. Before combining the two proteomic views, weights were scaled within each platform to prevent differences in weight magnitude between Olink and LC-MS/MS from determining the ranking. Bar colour represents signed significance, calculated as −log₁₀(P) multiplied by the direction of the normalized enrichment score. **G**, Heatmap of representative proteins contributing to the immune-response and oxidative-phosphorylation enrichments in Factor 1. Features were selected from the core-enrichment proteins of the corresponding GSEA terms. Within each term and proteomic platform, the five proteins with the largest directionally consistent scaled Factor 1 weights were retained. Rows show protein abundance standardised to a Z-score across samples. Columns represent individual patients and are ordered by TAPSE. Top annotations show TAPSE, RVIT, and RV state.

To analyse the two proteomic datasets jointly, we used Multi-Omics Factor Analysis (MOFA), which identifies latent molecular patterns based on coordinated variation across samples and estimates the contribution of each proteomic platform to these patterns. To avoid imposing a predefined number of factors, we compared models with increasing factor numbers and selected the final 11-factor model once additional factors consistently explained little additional variance (Extended Data Fig. 9H, Supplementary Data 8). We then tested the factor scores for associations with the haemodynamic and echocardiographic parameters. Factor 1 was associated with TAPSE and RVIT at FDR < 0.1 (Fig. 6C-D), and Factor 1 scores tended to be higher in patients with decompensated than compensated RV function, as defined by the TAPSE/sPAP ratio (P = 0.077; Fig. 6E). No factor was associated with mPAP, sPAP, PCWP or PVR. Separate differential abundance analyses of the LC-MS/MS and Olink datasets produced a similar pattern, with significant protein associations primarily observed for TAPSE and RVIT (Extended Data Fig. 10, Supplementary Data 9). Thus, proteomic variation was associated with RV systolic dysfunction, dilation and RV–pulmonary arterial uncoupling, rather than with the magnitude of pulmonary pressure load.

Gene set enrichment analysis of proteins ranked by their Factor 1 weights showed positive enrichment of adaptive immune responses, chemokine activity, inflammatory regulation and cell killing (Fig. 6F). Proteins contributing to these terms included mediators of innate immune and vesicular responses, such as VAMP8, CXCL8, LCP1 and ANXA1, together with lymphocyte- and antigen-presenting-cell-associated proteins including SH2D1A, CLEC10A and CLEC4G (Fig. 6G). Negative enrichment was dominated by mitochondrial respiration and oxidative phosphorylation. This included complex I components NDUFB7 and NDUFS6 and several cytochrome c oxidase subunits, including MT-CO2, COX4I1, COX5B and COX7A1.

### Shared transcriptomic and proteomic variation reflects RV fibrosis

We next integrated sample-level pseudobulk snRNA-seq profiles with the Olink and LC-MS/MS datasets from the 20 patients measured with all three approaches. This analysis was performed to determine which molecular patterns were shared across transcriptomic and proteomic layers. The final MOFA model contained seven factors (Fig. 7A, Extended Data Fig. 9I). Most factors captured variation predominantly from a single molecular layer or from the two proteomic layers. Factor 1 explained 40.8% of the transcriptomic variance but less than 1% of the variance in either proteomic dataset. Consistent with its predominantly transcriptomic origin, it was associated with mPAP, sPAP and PVR, recapitulating the pressure-load-associated pattern identified by the single-nucleus analyses (Fig. 7D). Factor 3 was largely specific to Olink, Factor 5 was predominantly driven by LC-MS/MS, and Factor 4 captured variation in both proteomic datasets with only a limited transcriptomic contribution. These findings indicate that the major sources of transcriptomic and proteomic variation in RV tissue were largely distinct.

Factor 2 was the clearest factor with substantial contributions from all three molecular layers, explaining 16.8% of the transcriptomic, 5.4% of the Olink and 11.8% of the LC-MS/MS variance (Fig. 7A). It was not associated with any of the tested haemodynamic or echocardiographic parameters but was strongly associated with histological RV fibrosis (FDR < 0.001; Fig. 7B-C). Selected features with high Factor 2 weights included immediate-early and AP-1-related genes (FOSB, FOS, JUNB, FOSL1, EGR3), profibrotic and TGF-β-associated genes (SERPINE1, TNC, CCN2, ADAMTS1, ITGA11, LTBP3), extracellular-matrix and adhesion components (VCAN, LUM, PRELP, FBLN1, MATN3, ITGB1, CD151), and genes related to myofibroblast differentiation and the contractile cytoskeleton (VIM, TAGLN, TPM4, FLNA, CNN1, ACTN1, VCL, MYH9) (Fig. 7E). These features originated from the transcriptomic, Olink and LC-MS/MS datasets, showing that fibrosis-related tissue remodelling was captured across all three molecular layers.

## Discussion

Right ventricular remodelling is a major determinant of outcome in pulmonary hypertension and advanced heart failure^1–3,8^. In PH due to left heart disease, elevated pulmonary pressure imposes a chronic load on the RV. The RV may initially compensate, but in some patients this response becomes insufficient and RV failure develops^5,6,8^. The molecular basis of this transition remains poorly defined in human tissue. To our knowledge, this is the first study to integrate single nucleus transcriptomics and tissue proteomics of the human RV in PH due to left heart disease. A major finding of our study was that transcriptional changes in the RV correlated with pulmonary pressure load rather than with the degree of RV decompensation.

Pathway analysis identified a transcriptional program overlapping with pathways previously implicated in RV adaptation and failure in human tissue and in animal models of PH and RV pressure overload. By linking these pathways to mPAP, our data position pulmonary pressure load as the likely trigger of a coordinated cardiomyocyte transcriptional response in the human RV. Mechanistically, this is consistent with increased wall stress activating mechanosensitive signalling, whereby cardiomyocytes and other resident cardiac cells sense elevated mechanical forces through mechanoreceptors and associated mechanotransduction pathways. The mPAP-associated program included genes involved in contractile and cytoskeletal remodelling, calcium handling, adrenergic signalling, autophagy, vesicle and endosomal transport, and glucose metabolism. Hypertrophic growth, sarcomeric remodelling and cytoskeletal adaptation are core features of the pressure-loaded RV in human and experimental studies^3,8,18^. In hypoxia-induced PH in calves, RV myocyte hypertrophy and selected hypertrophic transcripts increased with mean pulmonary arterial pressure, further supporting a direct link between pulmonary pressure load and hypertrophic remodelling^18^. The glucose metabolism signal is also consistent with the metabolic shift of the pressure-loaded RV, where increased glucose uptake, enhanced glycolytic activity and reduced reliance on oxidative metabolism have been described in human PH and experimental models^19–21^.

Autophagy and vesicle trafficking are more context-dependent. They may support adaptation by maintaining protein, membrane and organelle homeostasis under mechanical stress, but sustained or dysregulated autophagy may also become maladaptive^22–24^. In line with this, a recent multi-omics study of MCT-induced PH and pulmonary artery banding in rats identified changes in vesicle transport and autophagy at the protein, metabolite and transcript levels^25^. Although extracellular matrix remodelling is one of the most consistent features of RV remodelling in human PH and animal models^15,17,26,27^, higher mPAP in our cohort was associated with a fibroblast-centred TGF-β signalling network, but not with histological fibrosis or a transcriptomic signature of increased collagen deposition. Histological fibrosis was positively associated with the abundance of a distinct myofibroblast subset characterised by high expression of contractile genes. This suggests that pressure load can activate profibrotic signalling, but fibrosis is a complex, multifactorial process that may require additional inputs beyond mPAP^26,27^. In end-stage RV tissue, fibrosis may also reflect a cumulative remodelling process that developed before the terminal stage captured in our cohort. Studies in CTEPH suggest that part of the RV response to increased pressure is reversible: after pulmonary endarterectomy, haemodynamic unloading was associated with reduced RV transcriptional abnormalities, fibrosis and cardiomyocyte hypertrophy^17^.

However, fibrosis-related remodelling may regress less completely, as CMR studies after pulmonary endarterectomy show persistent extracellular volume expansion despite regression of cellular hypertrophy and improved RV stiffness^28^. These observations support our interpretation that the mPAP-associated pathways identified here represent a protective response aimed at preserving cardiomyocyte function under increased pressure load and mechanical stress.

These findings raise the question of why a pressure-responsive program preserves RV function in some patients but becomes insufficient in others. The distinction between compensated and decompensated RV states captures an important difference in RV function and prognosis. Clinical studies support this distinction, showing that reduced TAPSE/sPAP is associated with worse haemodynamics, functional class and outcome in PAH and heart failure, while invasive measures of RV–PA coupling also predict survival in PH^4,9,10^. An important implication of our findings is that this clinically defined transition to RV failure does not represent a transcriptionally distinct disease state. Garry et al. performed bulk RNA-seq on human RV tissue from patients with end-stage heart failure due to ischaemic or non-ischaemic cardiomyopathy undergoing transplantation^29^. Both compensated and failing RVs differed from unused donor hearts, but direct comparison of the failing and compensated groups identified only one significantly differentially expressed gene^29^. Similarly, Khassafi et al. showed that unsupervised transcriptomic clustering of clinically classified human RV samples identified molecular subgroups that only partially aligned with compensation status, with several clusters containing mixtures of normal, compensated, and decompensated RVs^14^. Progression from adaptive remodelling to overt RV decompensation may therefore arise not from activation of a decompensation-specific genetic program. Instead, this process is expected to involve the cumulative effects of post-transcriptional regulation, impaired protein homeostasis, metabolic remodelling, mitochondrial dysfunction, extracellular matrix remodelling, and structural alterations that are not fully captured at the transcriptomic level. Consistent with this concept, joint analysis of the LC-MS/MS and Olink datasets identified a protein abundance pattern associated with lower TAPSE, greater RV dilation and impaired RV–PA coupling, whereas no proteomic factor was associated with pulmonary haemodynamic parameters. This pattern combined higher abundance of proteins involved in immune and inflammatory processes with lower abundance of respiratory-chain proteins and other components of oxidative phosphorylation. Immune activation has been implicated in maladaptive RV remodelling in PH, including macrophage accumulation and NLRP3 inflammasome activation in decompensated RVs from patients with PAH and in monocrotaline and Sugen–hypoxia rat models^30^. In rats, failing Sugen–hypoxia RVs have likewise shown loss of mitochondrial proteins and impaired complex I respiration, changes that were not observed in functionally compensated, pressure-loaded RVs after pulmonary artery banding^31^. The comparatively modest associations observed in our cohort suggest that these immune and metabolic alterations are part of a broader combination of cellular, extracellular-matrix and biomechanical changes contributing to RV decompensation. When the snRNA-seq and proteomic datasets were integrated, most factors predominantly reflected variation in either RNA or protein abundance, indicating limited overlap in the biological patterns captured by the two molecular layers. The main shared factor was associated with histological fibrosis and contained canonical profibrotic, extracellular-matrix, adhesion and contractile genes and proteins. Fibrotic tissue remodelling therefore represented the clearest molecular process detected consistently at both the transcript and protein levels, although it was not associated with the haemodynamic or functional parameters examined.

Several limitations should be considered. The snRNA-seq analysis included 21 male patients with advanced DCM undergoing heart transplantation, limiting generalisability to women, earlier disease stages, and other causes of PH or RV failure. The overall sample size was limited, with small subgroups and missing haemodynamic data in some patients. Because human RV tissue was available only at transplantation, the study provides a cross-sectional view of advanced RV remodelling and lacks the temporal dimension needed to study the transition from compensated to decompensated states. We also lacked more detailed measures of RV performance and RV-pulmonary arterial coupling, such as RV ejection fraction, RV strain, or pressure-volume loop-derived indices of contractility and afterload. Finally, snRNA-seq and tissue proteomics capture only selected molecular layers and do not directly measure post-translational regulation, metabolic flux, or mitochondrial function.

In conclusion, pulmonary pressure load was the clinical variable most strongly associated with RV transcriptional remodelling in advanced DCM. Cardiomyocytes showed a broad pressure-load response involving contractile remodelling, calcium handling, autophagy, glucose metabolism and intracellular trafficking. Impaired RV-pulmonary arterial coupling, RV dysfunction and RV dilation did not define a separate cardiomyocyte transcriptional state, but were reflected more clearly, although partially, at the protein level, whereas fibrosis represented the main molecular process shared across transcriptomic and proteomic layers. Together, these findings indicate that pulmonary haemodynamic load, RV functional deterioration and myocardial fibrosis are associated with distinct molecular patterns, and support a model in which the transition to RV failure in advanced DCM reflects the insufficiency of pressure-load adaptation and downstream tissue-level remodelling, rather than a discrete transcriptional switch to decompensation.

## Methods

### Human samples and ethics approval

Right ventricular (RV) free-wall tissue was obtained from the explanted hearts of 44 patients with dilated cardiomyopathy (DCM) undergoing heart transplantation at the National Institute of Cardiology, Warsaw, Poland. Of these, 21 male patients were analysed by single-nucleus RNA sequencing (snRNA-seq), and all 44 patients were included in the LC-MS/MS and Olink Reveal proteomic analyses. All patients in the DCM group qualified for heart transplantation according to the criteria described in the 2021 ESC Guidelines for the diagnosis and treatment of acute and chronic HF^32^. The protocol was approved by the Local Ethics Committee (IK-NPIA-0021-34/1699/18 and IK-NPIA-0021-3.2062/24). The study was conducted in accordance with the Declaration of Helsinki and approved by Swissethics (BASEC-Nr. 2019-00058). Samples were snap-frozen in liquid nitrogen immediately after explantation and stored at -80 °C until processing.

### Clinical data and threshold definitions

Clinical, echocardiographic and haemodynamic data were obtained from medical records in a pseudonymised manner and are detailed in Supplementary Data 1. Echocardiographic variables included left ventricular ejection fraction (LVEF), left ventricular end-diastolic diameter (LVEDD), right ventricular inflow tract diameter (RVIT), and tricuspid annular plane systolic excursion (TAPSE). Haemodynamic variables obtained from right heart catheterisation included mean pulmonary arterial pressure (mPAP), systolic pulmonary arterial pressure (sPAP), pulmonary capillary wedge pressure (PCWP), and pulmonary vascular resistance (PVR). RV-pulmonary arterial coupling was estimated using the TAPSE/sPAP ratio.

For selected threshold-based transcriptomic analyses, patients were stratified according to markedly elevated pulmonary pressure load using mPAP ≥35 mmHg versus <35 mmHg, and RV state was defined as decompensated or compensated based on mPAP and TAPSE/sPAP. Patients with mPAP >20 mmHg and TAPSE/sPAP >0.36 were classified as compensated, whereas those with mPAP >20 mmHg and TAPSE/sPAP ≤0.36 were classified as decompensated. One patient with mPAP ≤20 mmHg was classified as not pressure overloaded. Patients with missing mPAP, or with mPAP >20 mmHg and missing TAPSE/sPAP, were not assigned to an RV state.

### Nuclei isolation and single-nuclei RNA sequencing

Snap-frozen RV tissue was cut into small pieces on ice using a sterile scalpel and transferred to a pre-chilled 2-ml Dounce homogenizer containing 500 µl of Nuclei EZ Lysis Buffer (Sigma-Aldrich). Tissue was homogenized with 10-15 strokes of the loose (A) pestle followed by 10-15 strokes of the tight (B) pestle. The homogenate was incubated on ice for 5 min and filtered through a 40-µm cell strainer. Nuclei were pelleted by centrifugation at 500g for 5 min at 4 °C, resuspended in 1 ml of lysis buffer, incubated on ice for 5 min, and centrifuged again under the same conditions. The pellet was washed in 1 ml of resuspension buffer (1× PBS, 1.0% BSA, 0.2 U/µl Ambion RNase inhibitor [Invitrogen]), centrifuged, and resuspended once more in 1 ml of the same buffer. After a final centrifugation, nuclei were resuspended in 500 µl of resuspension buffer supplemented with DAPI (10 µg/ml) and passed through a 35-µm filter cap for flow cytometry tubes. DAPI-positive nuclei were sorted on a BD FACSAria III cell sorter using a 100-µm nozzle. Nuclei integrity and morphology were verified by fluorescence microscopy before and after sorting. After sorting, nuclei were counted by trypan blue staining. Sorted nuclei were processed on the same day for library preparation.

Single-nucleus suspensions were loaded on a Chromium Next GEM Chip G (10x Genomics) targeting recovery of 10,000 nuclei per sample. Libraries were prepared with the Chromium Next GEM Single Cell 3′ v3.1 kit following the manufacturer’s protocol. Library quality and concentration were assessed with an Agilent Bioanalyzer (High Sensitivity DNA kit) or TapeStation (High Sensitivity D5000 ScreenTape).

Sequencing was performed at the Functional Genomics Center Zurich on an Illumina NovaSeq 6000 or NovaSeq X Plus platform, with a target depth of >40,000 reads per nucleus.

### Mapping, quality control and clustering

Raw sequencing reads were aligned to the GRCh38 human reference genome using Cell Ranger v7.0.0 (10x Genomics) with default settings. Ambient RNA contamination was corrected with CellBender v0.3.0 using default parameters. All subsequent analyses were performed in R v4.5.2.

For each sample, count matrices were converted into Seurat v4.4.0 objects. Doublets were identified with the scDblFinder package and removed. Nuclei were filtered to retain those with >500 detected genes, <1.5% mitochondrial reads, and 1,000-100,000 UMI counts.

Each sample was normalized and scaled with SCTransform, regressing out the percentage of mitochondrial and ribosomal reads. 3,000 variable features were selected with SelectIntegrationFeatures, and samples were integrated using Harmony with 50 principal components (PCs). Dimensionality reduction was performed with UMAP on the top 30 Harmony-corrected PCs, followed by clustering with FindNeighbors and FindClusters at a resolution of 0.2. One low-quality cluster characterised by low RNA content and high mitochondrial gene expression was removed.

Cluster markers were identified with FindAllMarkers (Wilcoxon rank-sum test, min.pct = 0.5, log₂FC threshold = 0.5). Clusters were annotated based on canonical marker genes and comparison with previously published human cardiac snRNA-seq datasets. For higher-resolution subclustering of major cell populations (cardiomyocytes, fibroblasts, endothelial cells, mural cells, macrophages and lymphocytes), nuclei were re-normalized and re-integrated using 2,000 variable features. Markers for these subclusters were identified with FindAllMarkers (min.pct = 0.1, log₂FC threshold = 0.25).

### Variance partitioning

Variance partitioning was used to estimate the contribution of clinical and technical variables to variation in the snRNA-seq and proteomics datasets. For snRNA-seq, nuclei from all annotated cell types were aggregated into one sample-level pseudobulk expression profile per patient. Pseudobulk libraries with fewer than 50,000 total counts were excluded. Lowly expressed genes were removed using the filterByExpr function from edgeR. Count data were then transformed using voom precision weights, and variance fractions were estimated with the fitExtractVarPartModel function from the variancePartition package.

The snRNA-seq model included age, TAPSE, RVIT, LVEF, LVEDD, mPAP, PCWP, PVR, and pseudobulk library size. Missing clinical values were handled using pairwise estimation, allowing samples with available data for a given variable to contribute to the estimation of that variable rather than restricting the analysis to samples with complete data for all variables. For visualisation, the average fraction of variance explained by each variable was calculated across all retained genes.

Variance partitioning was also performed for the LC-MS/MS and Olink proteomics datasets using the same modelling framework. For LC-MS/MS, the model included age, mPAP, RVIT, TAPSE, LVEF, and LVEDD. For Olink, total protein concentration measured by BCA assay was included as an additional technical covariate.

### Pseudobulk differential expression analysis

Differential expression analysis was performed using a sample-level pseudobulk approach. For each analysed cell type, raw snRNA-seq counts were aggregated per patient, generating one pseudobulk expression profile per sample and cell type. Pseudobulk differential expression analysis was performed for cell types with sufficient representation across patients, including cardiomyocytes, fibroblasts, endothelial cells, pericytes, smooth muscle cells, macrophages, neuronal cells, T lymphocytes and endocardial cells.

For each analysed cell type, genes expressed in 1% or fewer nuclei were removed before pseudobulk filtering. Pseudobulk libraries with fewer than 50,000 total counts were excluded. Additional lowly expressed genes were removed using the filterByExpr function from edgeR. For each cell type and clinical variable, samples with available data for the tested variable were retained. The tested clinical variable was scaled before model fitting to allow comparison of log2 fold-change estimates across parameters.

Pseudobulk counts were normalized for library size, dispersion was estimated, and differential expression was tested using quasi-likelihood negative binomial generalized linear models in edgeR. Models were fitted with the glmQLFit function, including the tested clinical variable and age as covariates to adjust for age differences between patients. For each gene, the association with the tested clinical variable was assessed using glmQLFTest on the coefficient corresponding to that variable. The continuous variables tested included mPAP, sPAP, PCWP, PVR, TAPSE, RVIT, LVEF and LVEDD. For threshold-based analyses, the same pseudobulk framework was used across these cell types to compare mPAP ≥35 versus <35 mmHg and compensated versus decompensated RV state as defined above. P values were adjusted for multiple testing using the Benjamini-Hochberg false discovery rate method. Genes were considered significant using FDR < 0.05 and absolute log2 fold change > 0.5.

### Cell-type and cell-state differential abundance analysis

Differential abundance analysis was used to test whether clinical variables were associated with changes in the relative abundance of annotated cell populations. The analysis was performed for major cell types and for cell states identified after subclustering cardiomyocytes, fibroblasts, endothelial cells, mural cells, macrophages and lymphocytes. Cell-type and cell-state proportions were logit-transformed using the getTransformedProps function from the speckle package. For each clinical variable, samples with available data for that variable were retained. Associations between logit-transformed proportions and clinical variables were tested using linear models fitted with the lmFit function from limma. Models included the tested clinical variable and age as covariates to adjust for age differences between patients. For each cell type or cell state, the model tested whether its relative abundance changed in association with the clinical variable after adjustment for age. The continuous variables tested included mPAP, sPAP, PCWP, PVR, TAPSE, RVIT, LVEF and LVEDD. P values were adjusted for multiple testing using the Benjamini-Hochberg false discovery rate method. For visualisation, untransformed proportions were plotted against the corresponding clinical variable.

### Gene set enrichment analysis and activity inference

Gene set enrichment analysis was performed on ranked pseudobulk differential expression results. For each analysis, genes were ranked using a signed statistic calculated as log2 fold change multiplied by −log10(P value). Gene symbols were converted to Entrez IDs, and genes without Entrez ID annotation were removed. Gene set enrichment analysis was performed with clusterProfiler using Gene Ontology terms and KEGG pathways, with a minimum gene set size of 10 genes. Redundant Gene Ontology terms were reduced using the simplify function from clusterProfiler, with a Wang semantic similarity cutoff of 0.75. When multiple similar terms were identified, the term with the lowest adjusted P value was retained.

For cardiomyocytes, recurrent mPAP-associated enriched terms were grouped into five biological programs: cytoskeleton and sarcomere remodelling, autophagy, adrenergic signalling, glucose metabolism, and vesicle transport. Genes contributing to the enriched terms assigned to each program were used to calculate gene-set scores. For each program, expression values were scaled for each gene across patients and then averaged to obtain one score per patient. Associations between gene-set scores and mPAP were tested using linear regression.

Transcription factor and pathway activities associated with the cardiomyocyte mPAP signature were inferred with decoupleR. The ranked cardiomyocyte mPAP differential expression signature was used as input. Transcription factor activity was inferred using the human CollecTRI transcription factor-target network and the run_ulm method, requiring at least 10 target genes per regulator. Transcription factors were retained if they were expressed in more than 5% of cardiomyocyte nuclei. Pathway activity was inferred using the human PROGENy pathway model and the run_ulm method.

The same activity inference analysis was also applied to subclustered cell states. Normalized single-nucleus expression matrices were used as input instead of ranked pseudobulk signatures. Transcription factors expressed in 1% or fewer nuclei in the analysed compartment were excluded.

### Histological fibrosis quantification and fibrosis-associated analyses

Right ventricular tissue sections were stained with Masson’s trichrome using a standard staining protocol. Whole-slide brightfield images were acquired with a Zeiss Axio Scan.Z1 slide scanner using a 20x objective. Fibrosis was quantified in Orbit Image Analysis using a supervised pixel classification approach. A pixel classification model was manually trained to distinguish background, non-fibrotic tissue and fibrotic tissue. For each section, the percentage of fibrotic area was calculated as the number of pixels classified as fibrotic divided by the total number of non-background tissue pixels.

Histological fibrosis values were integrated with the snRNA-seq metadata. Associations between fibrosis and fibroblast cell-state abundance, fibrosis-associated pseudobulk differential expression in fibroblasts, and downstream gene set enrichment analysis were performed as described above, using fibrotic area as the tested variable and age as a covariate.

### Cell-cell communication analysis

Cell-cell communication changes associated with pulmonary pressure load were inferred using MultiNicheNet. Patients with available mPAP values were stratified into groups using a threshold of 35 mmHg. All annotated cell types were considered as potential sender and receiver populations. Cell types were retained when they had at least 50 nuclei in a sample. Genes were considered expressed in a cell type when they were detected in at least 5% of nuclei and in at least 50% of samples within the relevant group. Differential expression between mPAP≥35 and mPAP<35 samples was calculated for each cell type, including age as a covariate. Differentially expressed genes with |log2 fold change| > 0.5 and P < 0.05 were used as target gene sets for ligand activity estimation. Ligand-receptor interactions and ligand-target links were inferred using the ligand-receptor network and ligand-target matrix provided by MultiNicheNet. Ligand-receptor pairs were prioritised by MultiNicheNet by integrating ligand and receptor expression, sender and receiver cell-type abundance, differential expression in receiver cell types, ligand-target regulatory potential, cell-type specificity and database support. Prioritised interactions were summarized using chord diagrams and bubble heatmaps. For the intercellular regulatory network, ligand-receptor-target links were further filtered to retain high-ranking target genes with concordant ligand-target expression correlations. The resulting network was used to visualise candidate signalling axes associated with the mPAP≥35 group.

### Preparation of RV tissue lysates for LC-MS/MS and Olink proteomics

Frozen RV tissue samples were cryopulverized with a mortar and pestle while maintained on dry ice and cooled with liquid nitrogen. The resulting tissue powder was collected and stored at −20 °C. After 24 h, the powder was resuspended in 100-300 µl of T-PER Tissue Protein Extraction Reagent (Thermo Fisher Scientific, 78510) supplemented with cOmplete Mini EDTA-free Protease Inhibitor Cocktail (Roche, 11836153001). The buffer volume was adjusted according to tissue powder weight. Lysates were centrifuged at 10,000g for 10 min at 4 °C, and the supernatant was collected. Protein concentration was measured using the Pierce BCA Protein Assay Kit (Thermo Fisher Scientific, 23225). Lysates were used without further dilution for LC-MS/MS. For Olink analysis, lysates were diluted to 1 mg/ml, and protein concentration was measured again before shipment. Olink Reveal proteomics was performed by Novogene.

### LC-MS/MS proteomics sample preparation and data acquisition

RV tissue lysates were mixed with SDS-buffer (1% SDS, 50 mM dithiothreitol, 100 mM Tris-HCl, pH 8) in a 1:2 v/v ratio (lysate:buffer) and incubated for 10 min at 95°C and 2000 rpm. An additional quality control (QC) sample was prepared from randomly chosen pooled clinical samples. Resulting samples were processed using the FASP (Filter-Aided Sample Preparation) method^33^ in the Microcon 10 kDa Centrifugal Filters (Merck, MCPRT010) according to a previously described protocol^34^. Briefly, a sample volume corresponding to 200 µg of protein was applied to the filter and washed by centrifugation multiple times (10,000 x g, room temperature, 30 min) using UA buffer (8 M urea, 100 mM Tris-HCl, pH 8.5).

Iodoacetamide (55 mM in UA buffer) was added to the filters, and samples were incubated in the dark at room temperature for 20 minutes. Next, filters were further washed by centrifugation several times with UA buffer, followed by DB buffer (50 mM Tris-HCl, pH 8.5). Filters were transferred to fresh tubes, and samples were digested with trypsin (2 µg per sample) overnight at 37°C. Peptides were eluted from filters with DB buffer. The peptide concentration was measured spectrophotometrically and approximately 10 µg of peptides were desalted on the in-house prepared C18 Stage Tips^35^. Samples were concentrated in a SpeedVac centrifuge at 45°C and iRT peptide standards (Biognosys IRT Kit, Bruker, 1816351) were added to samples according to manufacturer’s instructions.

LC-MS/MS measurements were conducted on a ZenoTOF 7600 mass spectrometer coupled to the M5 MicroLC-TE system (Sciex). Approximately 1.5 µg of peptides were injected for a single measurement. Peptide separation was performed in the trap-elute mode on a HALO C18 column (0.5 x 50 mm, 2.7 μm particle size, 160 Å pore size) in a 20-min gradient: 0-1 min: 5-7% B, 1-8 min: 7-10% B, 8-16 min: 10-20% B, 16-20 min: 20-35% B, followed by washing and reequilibration of the column (buffer A: 0.1% formic acid in water, buffer B: 0.1% formic acid in acetonitrile). Data were acquired using a Zeno SWATH DIA method in positive ion mode. Sixty variable acquisition windows covering the range of 300-1000 m/z were established using swathTuner on previously acquired test DDA measurements of selected clinical samples^36^. The method included a survey scan in the range of 300-1600 m/z for 50 ms followed by a TOF MS/MS scan in each acquisition window in the range of 50-2000 Da for 10 ms. The total cycle time was 1.138 s. Samples were injected in random order with a blank measurement after every three sample measurements, and a QC measurement following every ten clinical samples.

Acquired raw data were processed in DIA-NN 2.3.2^37^ using a predicted spectral library built from the *Homo sapiens* proteome (UP000005640) downloaded from UniProt (March 15, 2026) supplemented with sequences of iRT peptides and cRAP protein sequences to monitor contamination. Cysteine carbamidomethylation was set as a fixed modification. The maximum number of variable modifications was set to 3. N-terminal acetylation and methionine oxidation were set as variable modifications, while the maximum number of missed cleavages was set to 2. Match-between-runs was enabled, and RT-dependent cross-run normalization was performed. Output was organized in protein groups and filtered at 1% FDR.

### Proteomics data preprocessing and differential abundance analysis

LC-MS/MS and Olink Reveal data were first processed and analysed separately. Protein abundance matrices were matched with sample metadata and clinical variables before downstream analysis. LC-MS/MS protein intensities were log2-transformed as log2(intensity + 1), whereas Olink analyses were performed using normalized protein expression (NPX) values. Differential protein abundance was tested separately for each clinical variable using linear models implemented in limma. Continuous clinical variables were rescaled before model fitting. For each model, samples with available values for the tested clinical variable and covariates were retained, and proteins with insufficient data for model fitting were excluded. Models included the tested clinical variable and age as covariate. For Olink analyses, total protein concentration measured by BCA assay was included as an additional covariate. P values were adjusted for multiple testing using the Benjamini-Hochberg false discovery rate method. Proteins with FDR < 0.2 were considered significant in the clinical association analyses.

### Multi-Omics Factor Analysis

Multi-Omics Factor Analysis was performed using MOFA2. Two models were fitted. The first model jointly analysed Olink and LC-MS/MS data from patients measured with both proteomic platforms. Before model fitting, the linear effect of sample protein concentration was removed from each Olink protein using the removeBatchEffect function in limma. Olink proteins with non-zero variance after correction were retained. For LC-MS/MS, proteins detected in at least 50% of samples and with non-zero variance were retained.

Missing LC-MS/MS values were not imputed and were handled as missing during model fitting.

The second model integrated sample-level pseudobulk snRNA-seq, Olink and LC-MS/MS data from the 20 patients measured with all three approaches. For the RNA dataset, genes detected in at least 1% of nuclei from patients with DCM were retained, and mitochondrial and ribosomal genes were excluded.

Counts were aggregated across all nuclei from each patient and transformed using the variance-stabilizing transformation implemented in DESeq2. The 2,000 genes with the highest variance across patients were selected.

All input datasets contained continuous transformed values and were modelled using Gaussian likelihoods. Each dataset was scaled to unit variance during model fitting. Clinical and histological variables were not included during model training. Models containing 5–15 factors were fitted using three random seeds. For each factor number, the run with the highest evidence lower bound was retained. The final model was selected immediately before the first of three consecutive models in which the weakest factor explained less than 2% of the combined variance across datasets. This resulted in 11 factors for the proteomics-only model and seven factors for the model integrating RNA, Olink and LC-MS/MS data.

Associations between factor scores and mPAP, sPAP, PCWP, PVR, TAPSE, RVIT, LVEF, and LVEDD were tested using separate linear models. Factor score was used as the outcome and the scaled clinical parameter as the predictor, adjusting for scaled age. P values were adjusted across all factor-parameter combinations using the Benjamini–Hochberg method, and associations with FDR <0.10 were considered significant. Differences in factor scores between compensated and decompensated patients were tested using linear models adjusted for age. Associations between factor scores and histological fibrosis were tested in the model integrating all three molecular datasets using linear models adjusted for age. Fibrotic area was scaled before model fitting, and P values were adjusted across factors using the Benjamini– Hochberg method. The relationship between Factor 2 scores and fibrotic area was additionally assessed using Spearman correlation. Gene Ontology gene set enrichment analysis was performed with clusterProfiler as described above, using proteins ranked by their Factor 1 weights. To combine the Olink and LC-MS/MS results, weights were scaled within each platform by the largest absolute weight. Scaled weights were averaged for proteins measured by both platforms. Gene sets containing 5–500 genes were tested. Representative proteins shown in the heatmap were selected from the core-enrichment features of the immune-response and oxidative-phosphorylation terms. Up to five proteins per term and proteomic platform with the largest weights in the direction of enrichment were retained. For the fibrosis-associated Factor 2, weights were oriented in the direction of increasing histological fibrosis. Selected features with high fibrosis-oriented weights from the RNA, Olink and LC-MS/MS datasets were annotated according to their biological functions and displayed across patients ordered by fibrotic area.

## Data availability

The raw snRNA-seq data and the processed Seurat object have been deposited in ArrayExpress under accession number E-MTAB-17557. The mass spectrometry proteomics data have been deposited to the ProteomeXchange Consortium via the PRIDE partner repository with the dataset identifier PXD082786. The Olink Reveal NPX data are provided as Supplementary Data 7.

## Code availability

All code used for data analysis is publicly available on GitHub at https://github.com/GinoBonazza/DCM_snRNAseq. The analyses were implemented using the workflowr package, which provides version control and generates HTML reports containing the code, results and figures. The reports are available through the project website at https://ginobonazza.github.io/DCM_snRNAseq/.

## Author contributions

G.A.B. processed tissue samples for snRNA-seq and proteomic analyses, prepared libraries for snRNA-seq, performed the downstream bioinformatic and statistical analyses, developed the analysis code, contributed to the study design and analytical strategy, prepared the figures and wrote the manuscript. A.P. and M.M. provided the tissue samples and clinical data. M.M. also contributed to interpretation of the results and revision of the manuscript. A.B. performed LC-MS/MS sample preparation, data acquisition and initial data processing. W.Ś.-R. assisted with LC-MS/MS data acquisition and instrumentation. A.L. performed Masson’s trichrome staining. P.L. facilitated access to the transplant cohort, tissue samples and associated clinical data. O.D. provided feedback on the study. P.B. and G.K. conceived and designed the study, supervised the project, contributed to interpretation of the results and revised the manuscript. All authors reviewed and approved the final manuscript.

## Acknowledgements

We thank Cristian Iperi for guidance on MOFA analysis; Mark Robinson and Christof Seiler for helpful discussions and feedback on data analysis; Elena Pachera for guidance on snRNA-seq library preparation; and Magdalena Czepiec and Anna Dobosz for assistance with sample processing for the proteomic analyses.

## Extended Data figures

**Extended Data Fig. 1:**
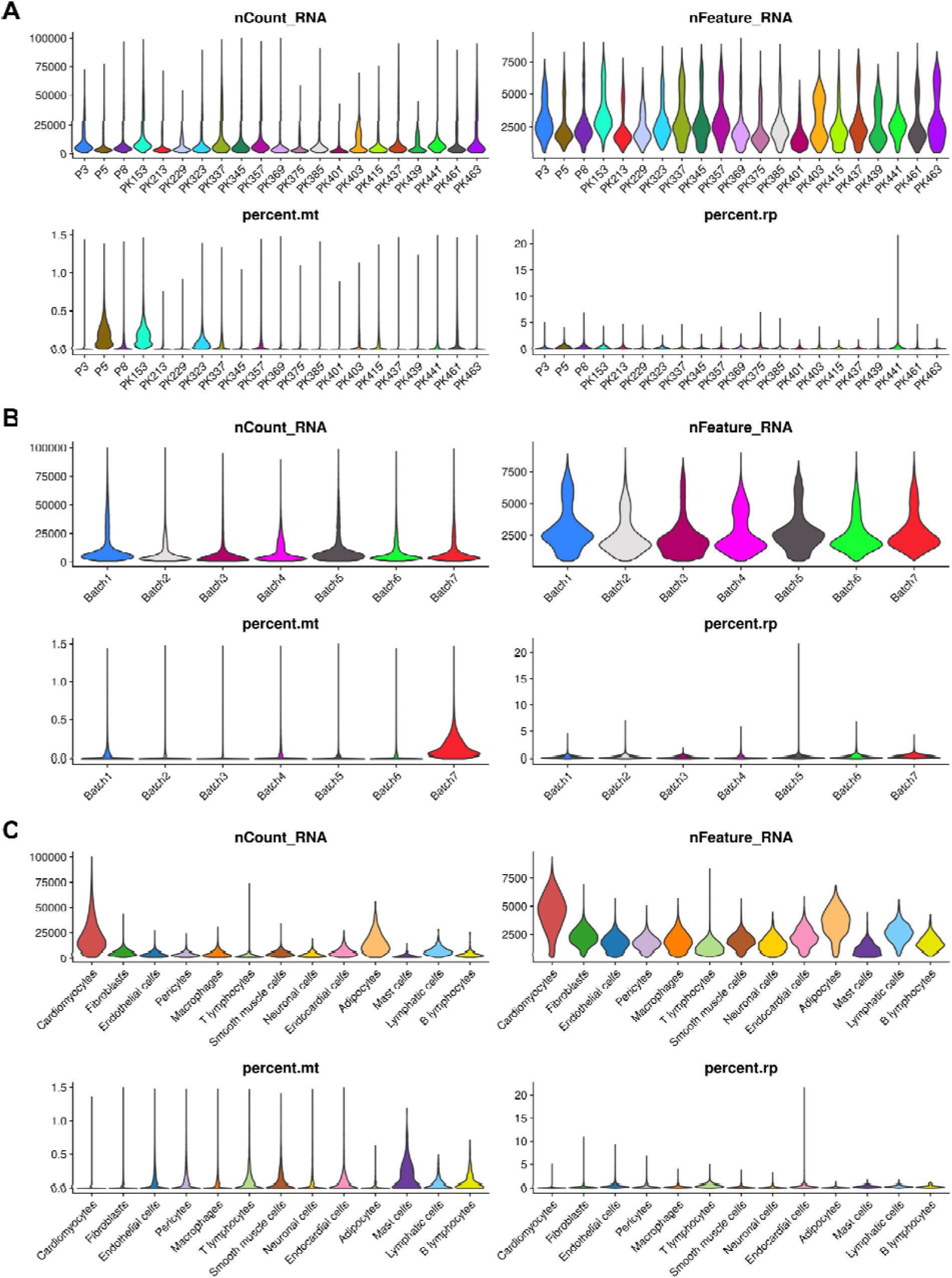
Quality control metrics of the final snRNA-seq dataset. **A**, Violin plots showing the distribution of quality control metrics across individual DCM samples after removal of low-quality nuclei, including total RNA counts per nucleus, number of detected genes per nucleus, percentage of mitochondrial transcripts, and percentage of ribosomal transcripts. **B**, Distribution of the same quality control metrics across sequencing or processing batches. Up to four samples were processed in the same batch. **C**, Distribution of quality control metrics across annotated cell types in the final integrated dataset. After quality control, the final dataset contained 136,769 nuclei and 30,376 detected genes. Final cell type annotations included cardiomyocytes, fibroblasts, endothelial cells, pericytes, macrophages, T lymphocytes, smooth muscle cells, neuronal cells, endocardial cells, adipocytes, mast cells, lymphatic cells, and B lymphocytes.

**Extended Data Fig. 2:**
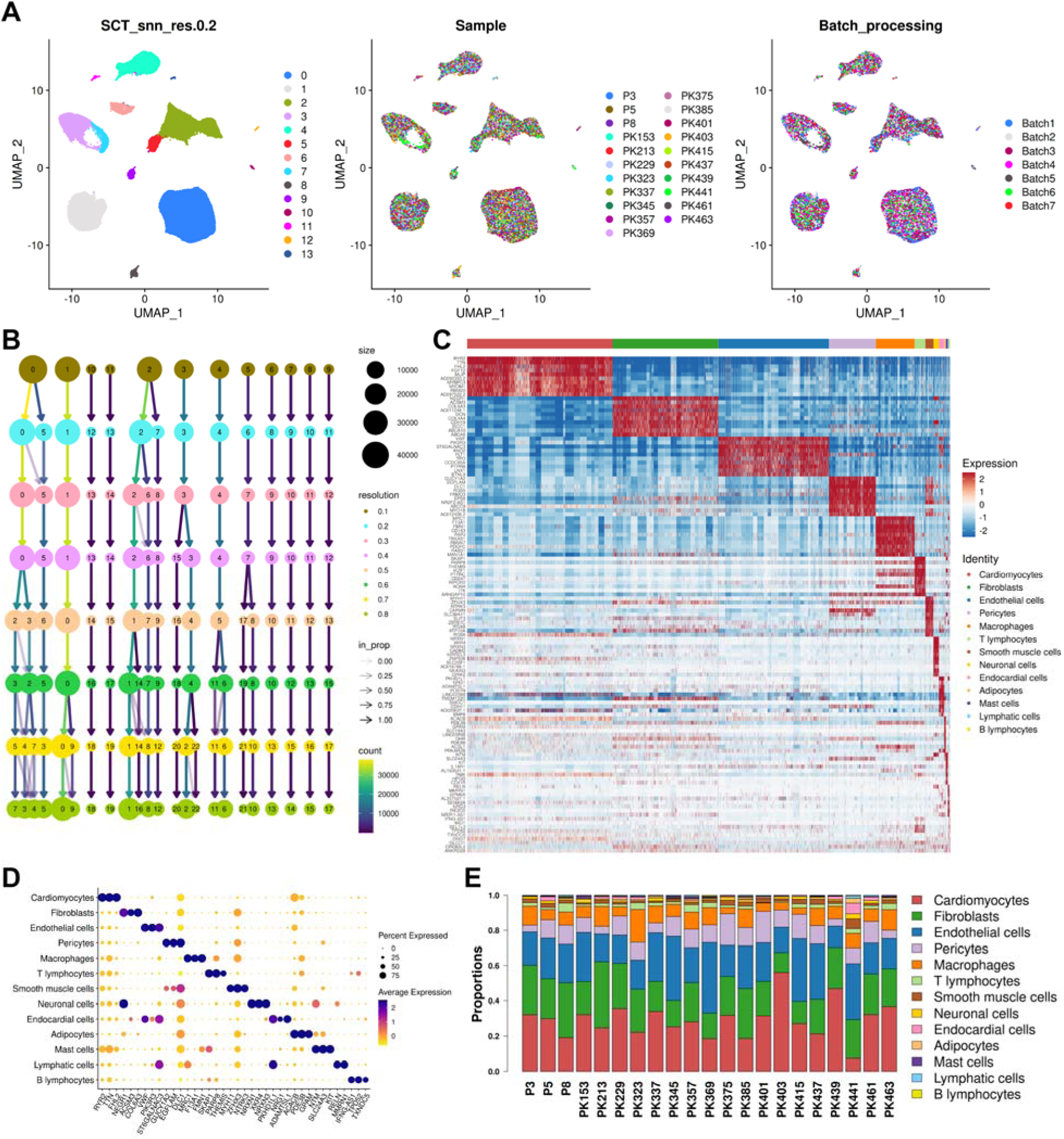
Integration, clustering, and annotation of the final snRNA-seq dataset. **A**, UMAP representation of the final integrated DCM snRNA-seq dataset, shown by unsupervised clusters at the selected clustering resolution of 0.2, by sample, and by processing batch. Integration across samples and batches showed good mixing of nuclei. A low clustering resolution was used to identify major cell populations, resulting in 14 clusters. Clusters 2 and 5 were both annotated as endothelial cells and were merged, giving 13 final major cell types. **B**, Clustering tree showing the progressive split of clusters across increasing clustering resolutions. The tree indicates a stable clustering structure, with limited reassignment of nuclei between clusters across resolutions. **C**, Heatmap showing the top 10 marker genes for each annotated major cell type. Columns represent nuclei grouped by cell type, and rows represent marker genes. **D**, Dot plot displaying the characteristic marker genes for each cell population. The top three marker genes by log₂ fold change (log₂FC) are shown for each population. **E**, Stacked bar plot showing the proportion of each annotated cell type in each DCM sample.

**Extended Data Fig. 3:**
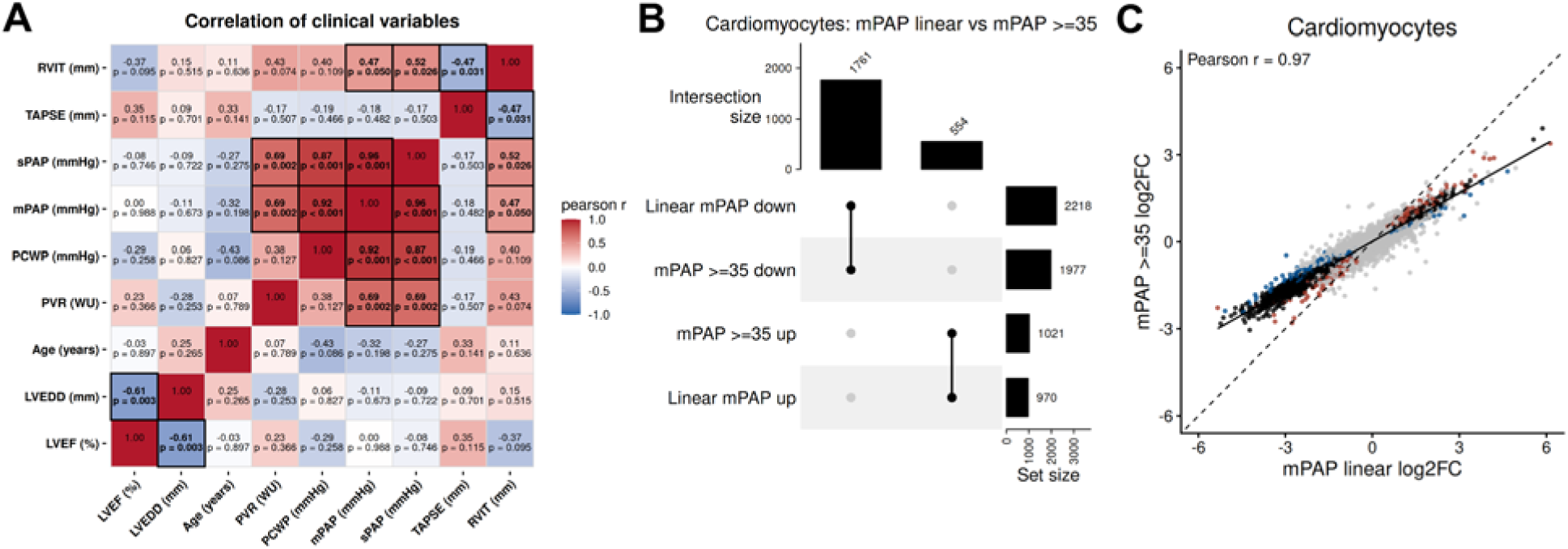
Clinical covariates and comparison of continuous and threshold-based mPAP differential expression. **A**, Correlation matrix of the clinical variables used for differential expression analysis. Pulmonary hypertension related parameters, including mPAP, sPAP, PCWP, and PVR, were strongly correlated with each other. TAPSE and RVIT showed a negative correlation, consistent with the relationship between RV function and RV dilation. LVEF and LVEDD showed a similar inverse relationship. Age did not show strong correlations with the main clinical parameters. **B**, UpSet plot showing the overlap between cardiomyocyte genes associated with mPAP in the continuous model and genes differentially expressed between patients with mPAP ≥35 mmHg and mPAP <35 mmHg. Overlaps are shown separately for genes with increased and decreased expression. **C,** Scatter plot comparing log₂FC estimates from the continuous mPAP model and the mPAP ≥35 mmHg group comparison in cardiomyocytes. The strong correlation between both models indicates that the threshold-based comparison captures the same mPAP-associated transcriptional signal observed in the continuous analysis.

**Extended Data Fig. 4:**
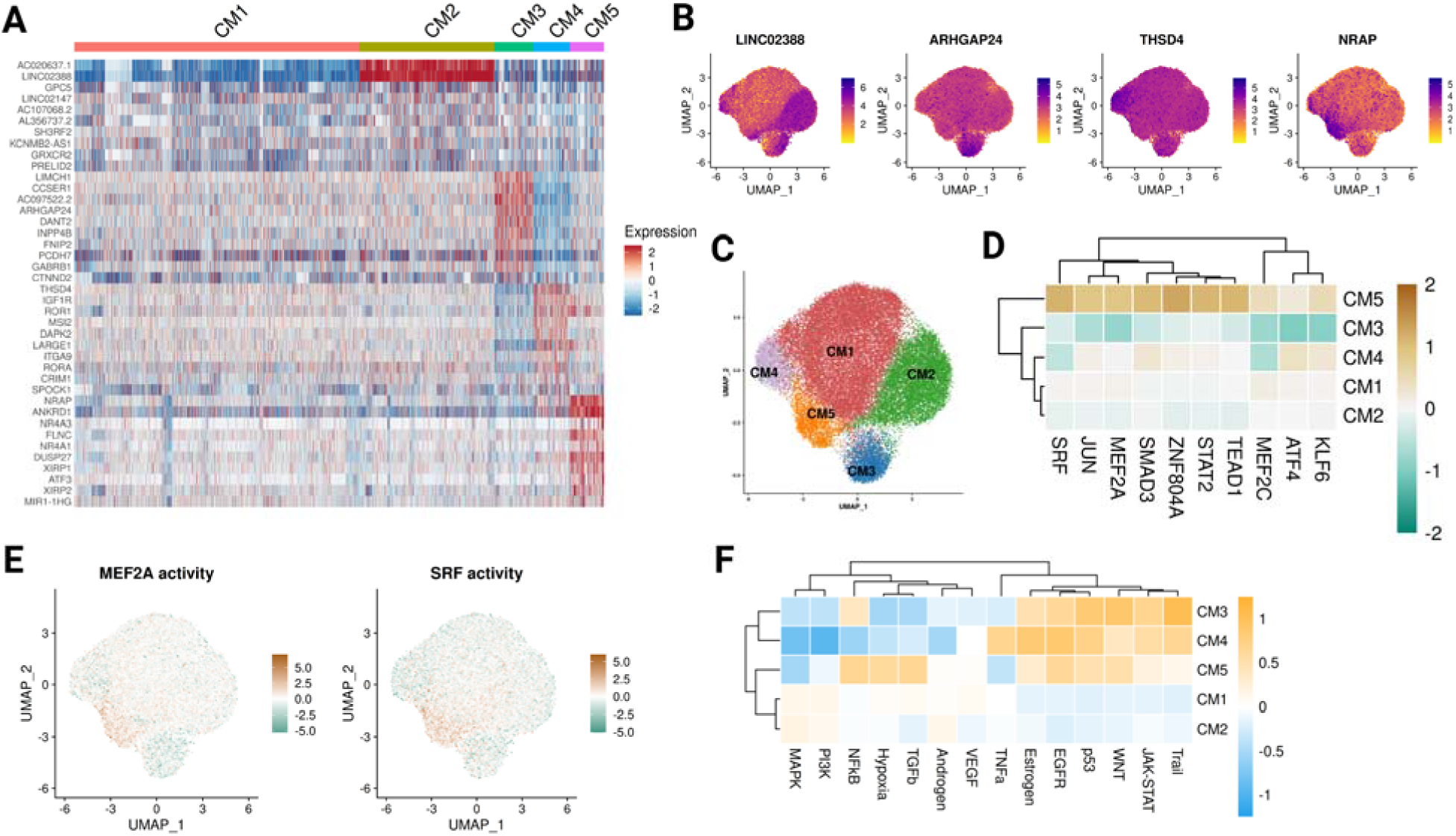
Subset analysis of cardiomyocytes. **A**, Heatmap showing the top 10 marker genes for each cardiomyocyte subset. No markers were detected for CM1. Columns represent nuclei grouped by subset, and rows represent marker genes. **B**, UMAP feature plots showing the expression of selected marker genes across cardiomyocyte subsets. **C**, UMAP representation of the five cardiomyocyte subsets (CM1–CM5). **D**, Heatmap of inferred transcription factor activity across cardiomyocyte subsets. CM5 showed higher predicted activity of several transcription factors, including MEF2A and SRF. **E**, UMAP representation of inferred MEF2A and SRF activity in cardiomyocytes. **F**, Heatmap of inferred pathway activity across cardiomyocyte subsets.

**Extended Data Fig. 5:**
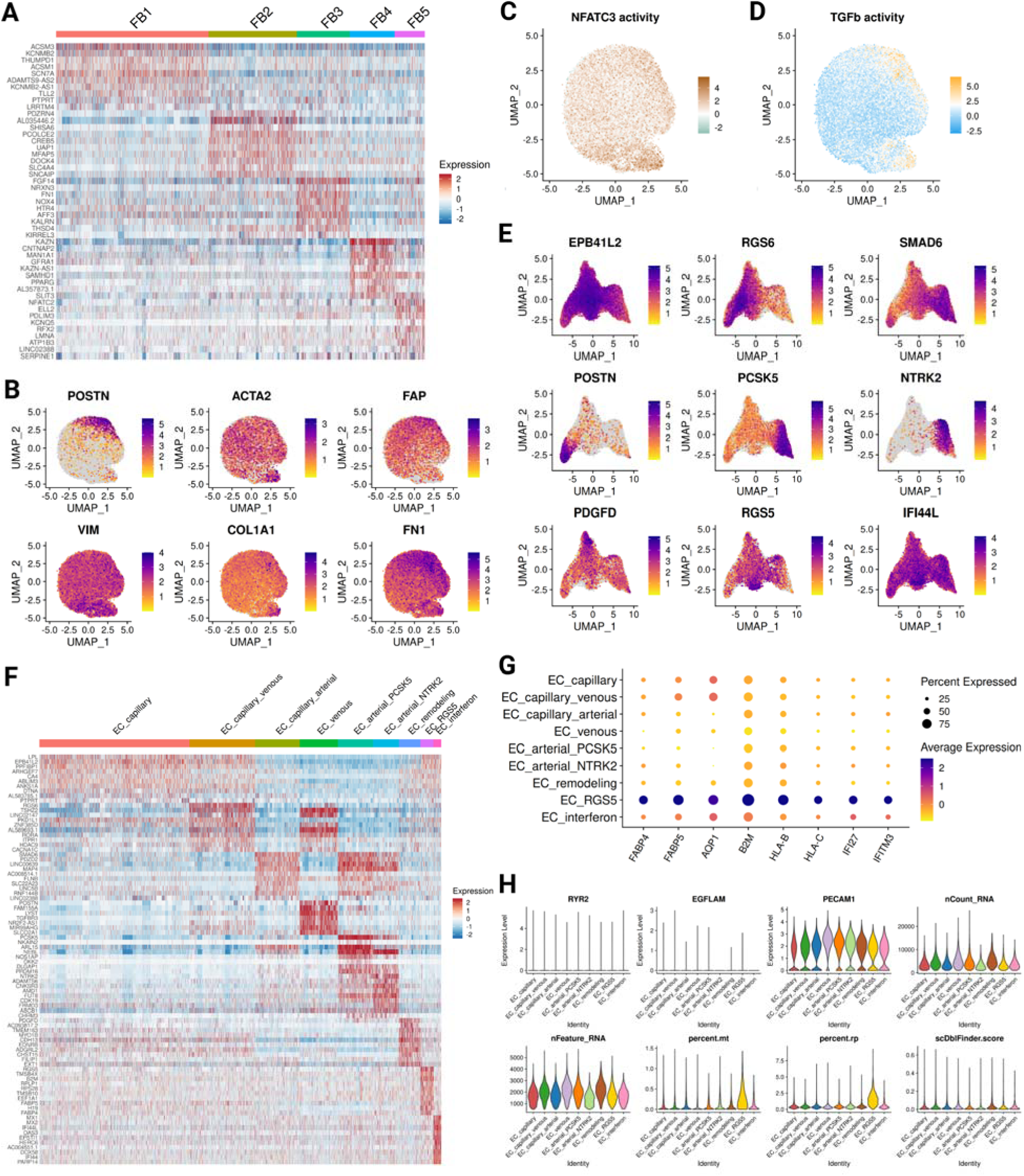
Subset analysis of fibroblasts and endothelial cells. **A**, Heatmap showing the top 10 marker genes for each fibroblast subset. Columns represent nuclei grouped by subset, and rows represent marker genes. **B**, UMAP feature plots showing selected fibroblast marker genes related to fibroblast activation, myofibroblast phenotype, and extracellular matrix remodelling. **C**, UMAP representation of inferred NFATC3 activity in fibroblasts, showing higher activity in FB5. **D**, UMAP representation of inferred TGFβ pathway activity in fibroblasts, showing higher activity in FB3 and FB5. **E**, UMAP feature plots showing selected marker genes across endothelial cell subsets. **F**, Heatmap showing the top 10 marker genes for each endothelial cell subset. Columns represent nuclei grouped by subset, and rows represent marker genes. **G**, Dot plot showing selected marker genes used to characterise the EC_RGS5 subset. Dot size indicates the fraction of nuclei expressing each gene and colour indicates average scaled expression. **H**, Violin plots showing selected quality control features and expression of selected marker genes across endothelial cell subsets, including expression of RYR2, EGFLAM, and PECAM1, as well as nCount_RNA, nFeature_RNA, percent.mt, percent.rp, and scDblFinder score. The EC_RGS5 subset showed higher mitochondrial and ribosomal transcript percentages and expressed RGS5, a marker commonly associated with pericytes. However, this subset did not show high expression of EGFLAM, one of the top pericyte markers, did not have increased doublet scores, and retained high expression of the endothelial marker PECAM1. Together, these features support the interpretation that EC_RGS5 represents an endothelial subset with RGS5 expression rather than pericyte contamination or doublets.

**Extended Data Fig. 6:**
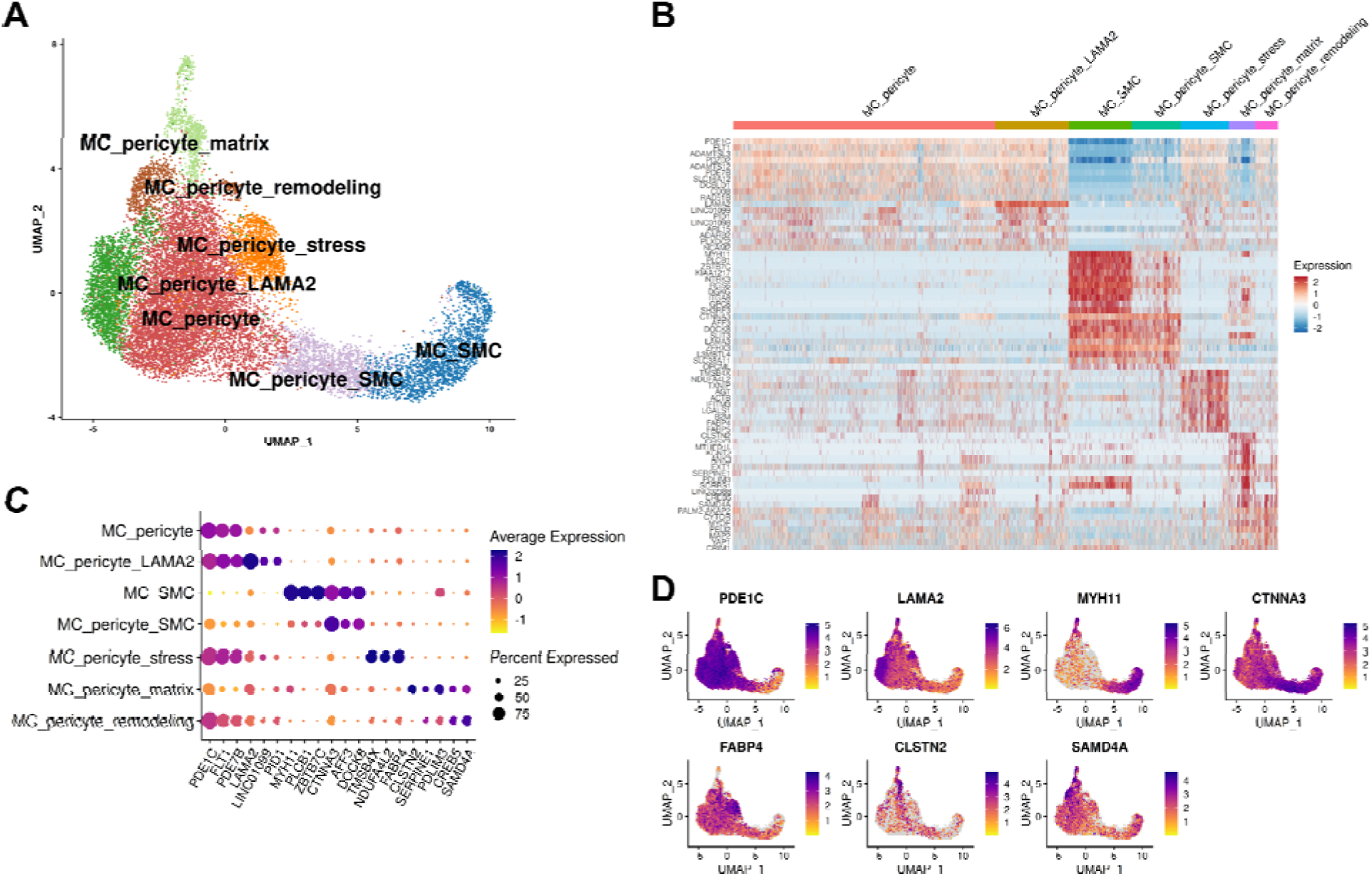
Subset analysis of mural cells. **A**, UMAP representation of mural cells, showing the seven annotated pericyte and smooth muscle cell states. **B**, Heatmap showing the top 10 marker genes for each mural cell subset. Columns represent nuclei grouped by subset, and rows represent marker genes. **C**, Dot plot summarising the top three marker genes for each mural cell subset. Dot size indicates the fraction of nuclei expressing each gene and colour indicates average scaled expression. **D**, UMAP feature plots showing the expression of selected marker genes across mural cell subsets.

**Extended Data Fig. 7:**
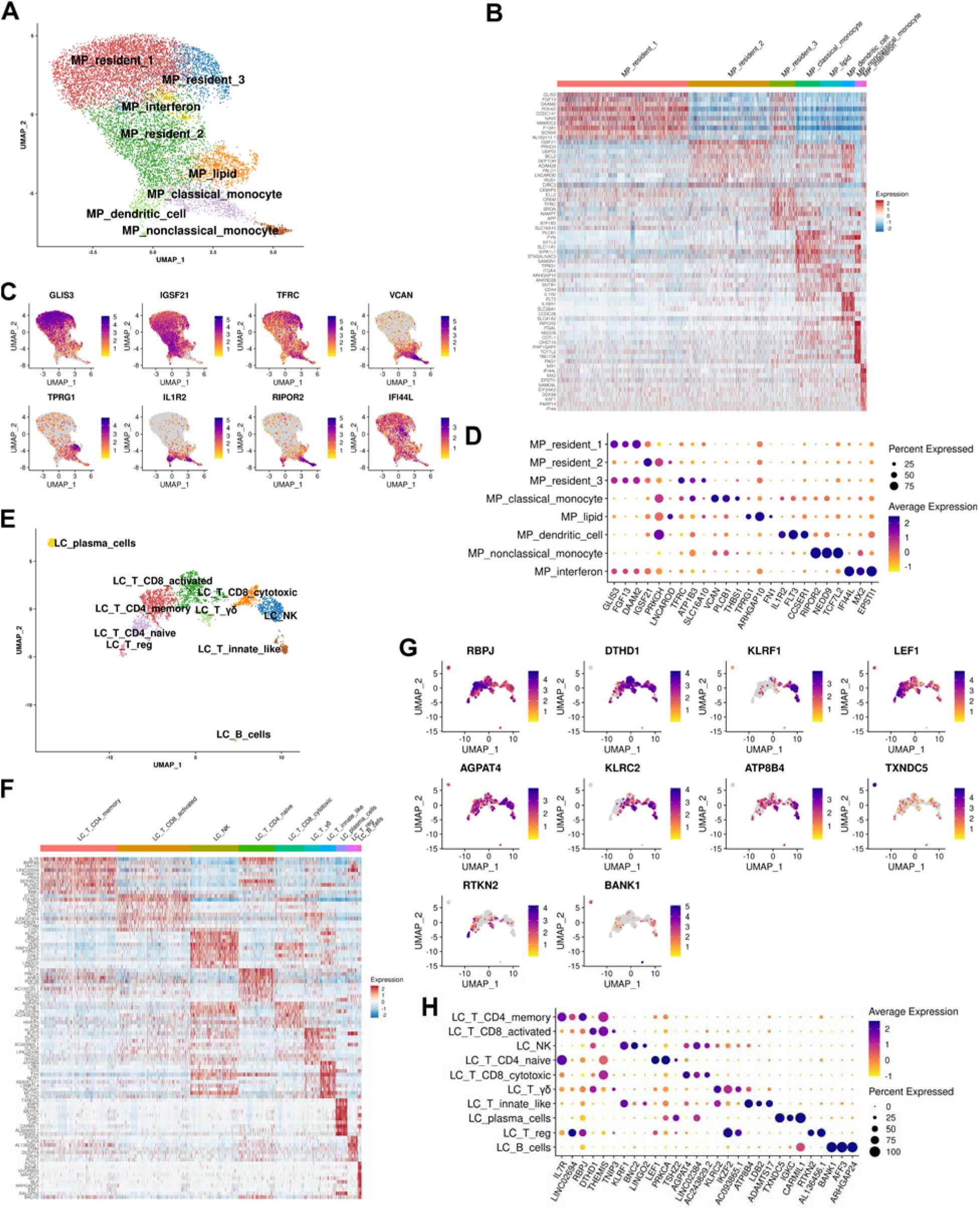
Subset analysis of immune cells. **A**, UMAP representation of macrophages, showing the annotated macrophage subsets. **B**, Heatmap showing the top 10 marker genes for each macrophage subset. Columns represent nuclei grouped by subset, and rows represent marker genes. **C**, UMAP feature plots showing the expression of selected marker genes across macrophage subsets. **D**, Dot plot summarising the top three marker genes for each macrophage subset. Dot size indicates the fraction of nuclei expressing each gene and colour indicates average scaled expression. **E**, UMAP representation of lymphocytes, showing the annotated lymphocyte subsets. **F**, Heatmap showing the top 10 marker genes for each lymphocyte subset. Columns represent nuclei grouped by subset, and rows represent marker genes. **G**, UMAP feature plots showing the expression of selected marker genes across lymphocyte subsets. **H**, Dot plot summarising the top three marker genes for each lymphocyte subset. Dot size indicates the fraction of nuclei expressing each gene and colour indicates average scaled expression.

**Extended Data Fig. 8:**
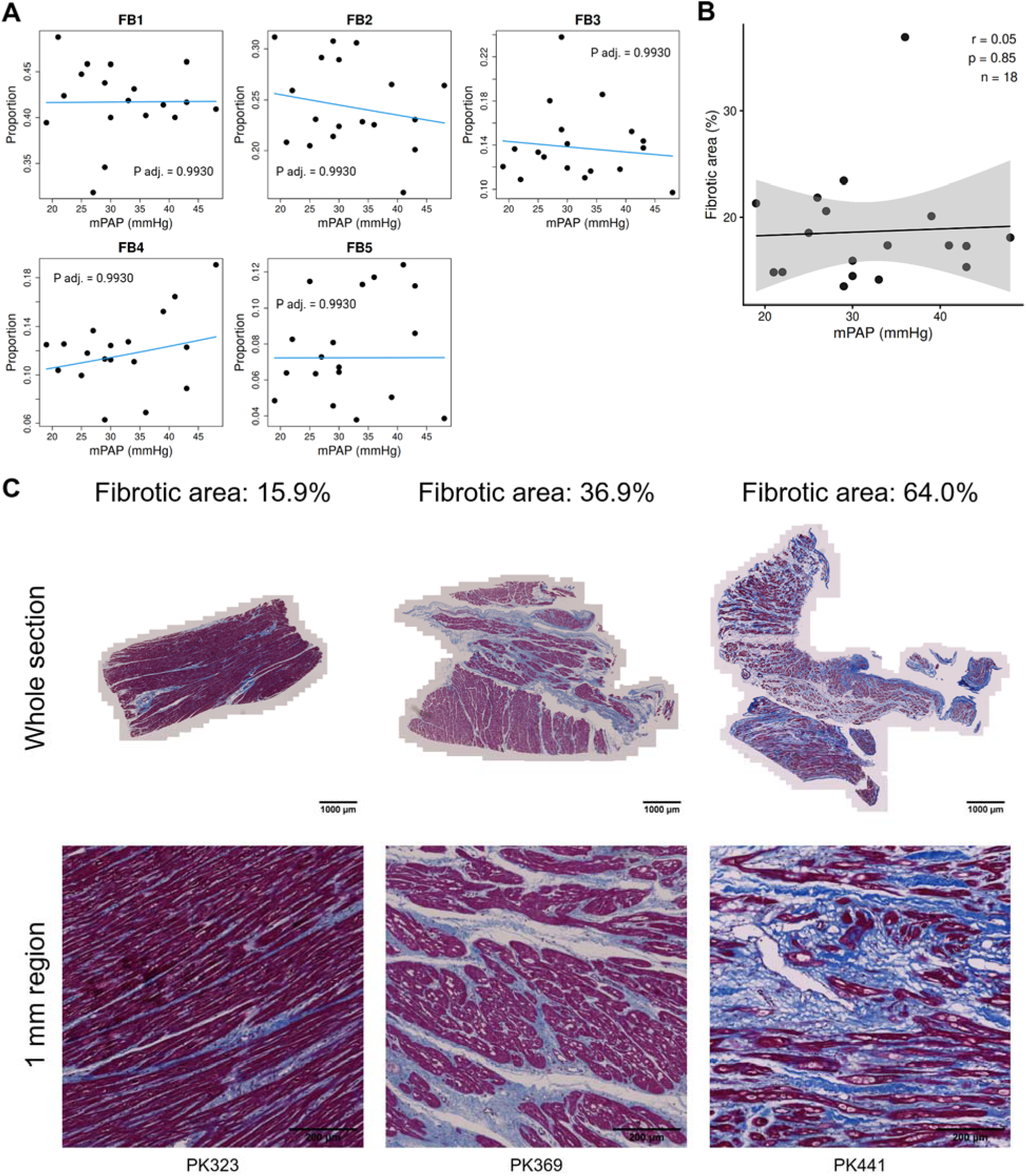
Fibroblast subset abundance and fibrosis in relation to mPAP. **A**, Association between fibroblast subset proportions and mPAP across DCM samples. Differential abundance was tested using the propeller method on logit-transformed proportions with a limma empirical Bayes linear model, including age as a covariate. **B**, Correlation between histological fibrosis, measured as fibrotic area (%), and mPAP across DCM samples. Pearson correlation showed no significant association between fibrosis and mPAP. **C**, Representative Masson’s trichrome-stained RV sections from three patients spanning increasing levels of histological fibrosis (15.9%, 36.9% and 64.0% fibrotic area). Whole-section images are shown above and corresponding higher-magnification regions below. Scale bars are indicated in the images.

**Extended Data Fig. 9:**
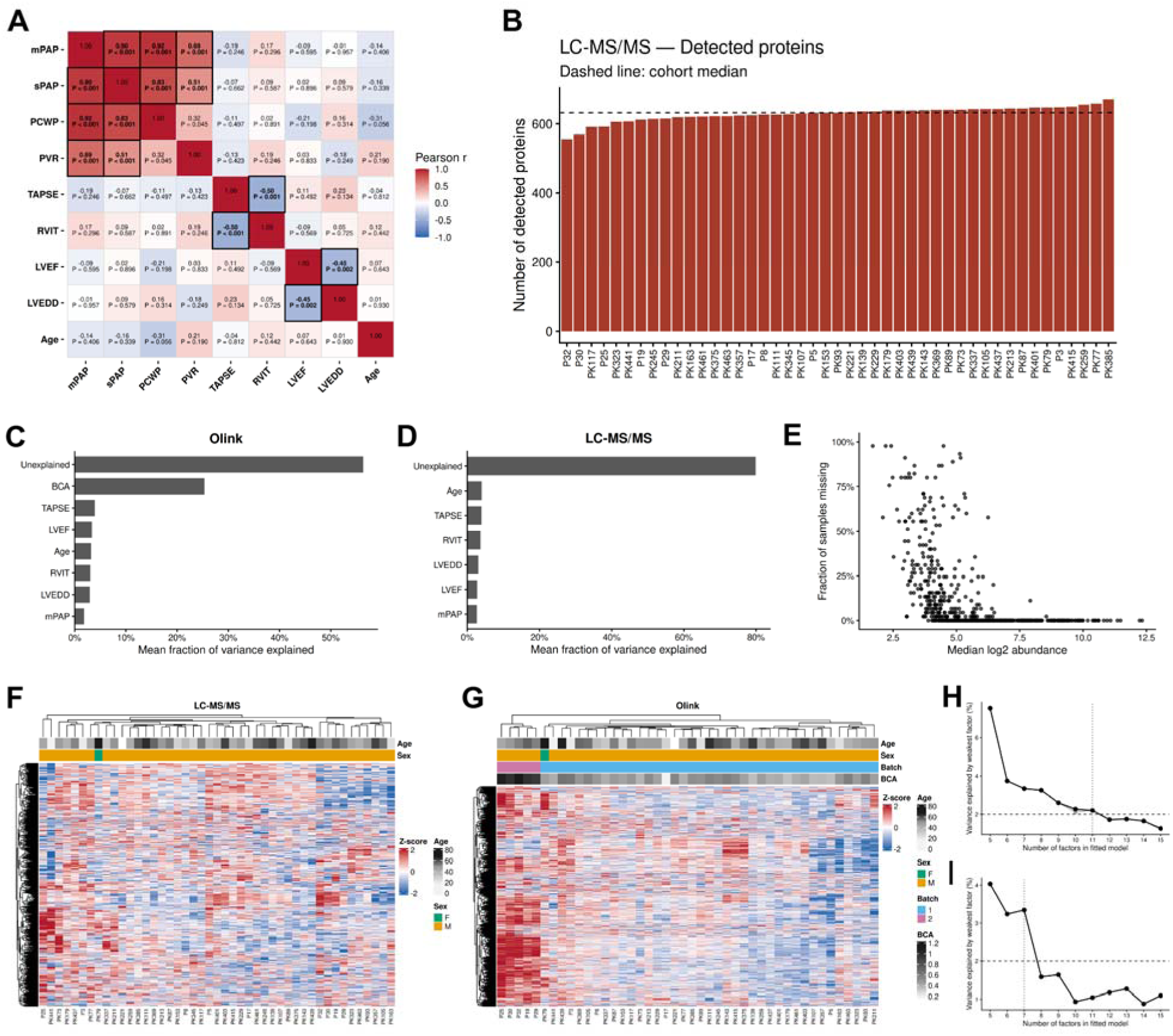
Quality assessment and model selection for the proteomic and multi-omics analyses. **A**, Correlation matrix of the clinical variables in the proteomics cohort. Pulmonary hypertension related parameters, including mPAP, sPAP, PCWP, and PVR, were strongly correlated with each other. TAPSE and RVIT showed a negative correlation, consistent with the relationship between RV function and RV remodelling. LVEF and LVEDD showed a similar inverse relationship. Age did not show strong correlations with the main clinical parameters. **B**, Bar plot showing the number of detected proteins in each sample in the final LC-MS/MS dataset after exclusion of low-quality samples. The number of detected proteins was comparable across samples. **C-D**, Variance partitioning analysis of the Olink (**C**) and LC-MS/MS (**D**) datasets. In the Olink dataset, sample protein concentration measured by BCA assay explained a substantial fraction of the variance and was therefore included as a covariate in downstream differential abundance analyses. In both datasets, clinical variables explained only a small fraction of the variance, in contrast to the snRNA-seq data, and most variance remained unexplained. **E**, Scatter plot showing, for each LC-MS/MS protein, the relationship between the fraction of missing values and the median log2 abundance. Proteins with lower abundance tended to have a higher fraction of missing values, whereas more abundant proteins were detected more consistently across samples. **F**, Heatmap showing scaled abundance values for all proteins in the LC-MS/MS dataset across samples. No major outlier samples were observed. **G**, Heatmap showing scaled abundance values for all proteins in the Olink dataset across samples. Samples P25, P39, P32, P19, and P29 were processed in a second batch and showed globally higher protein abundance, consistent with higher sample concentration. This batch-related effect was accounted for in downstream analyses. **H–I**, Selection of the number of factors for the proteomics-only MOFA model (**H**) and the model integrating pseudobulk snRNA-seq, Olink and LC-MS/MS data (**I**). Models containing 5–15 factors were fitted using three random seeds. For each number of factors, the run with the highest evidence lower bound was retained. Points show the combined percentage of variance explained by the weakest factor across all views. The horizontal dashed line indicates the prespecified 2% threshold. The selected model was the model immediately preceding the first of three consecutive models in which the weakest factor explained less than 2% of the combined variance. Vertical dotted lines indicate the selected models, containing 11 factors for the proteomics-only analysis and seven factors for the three-view analysis.

**Extended Data Fig. 10:**
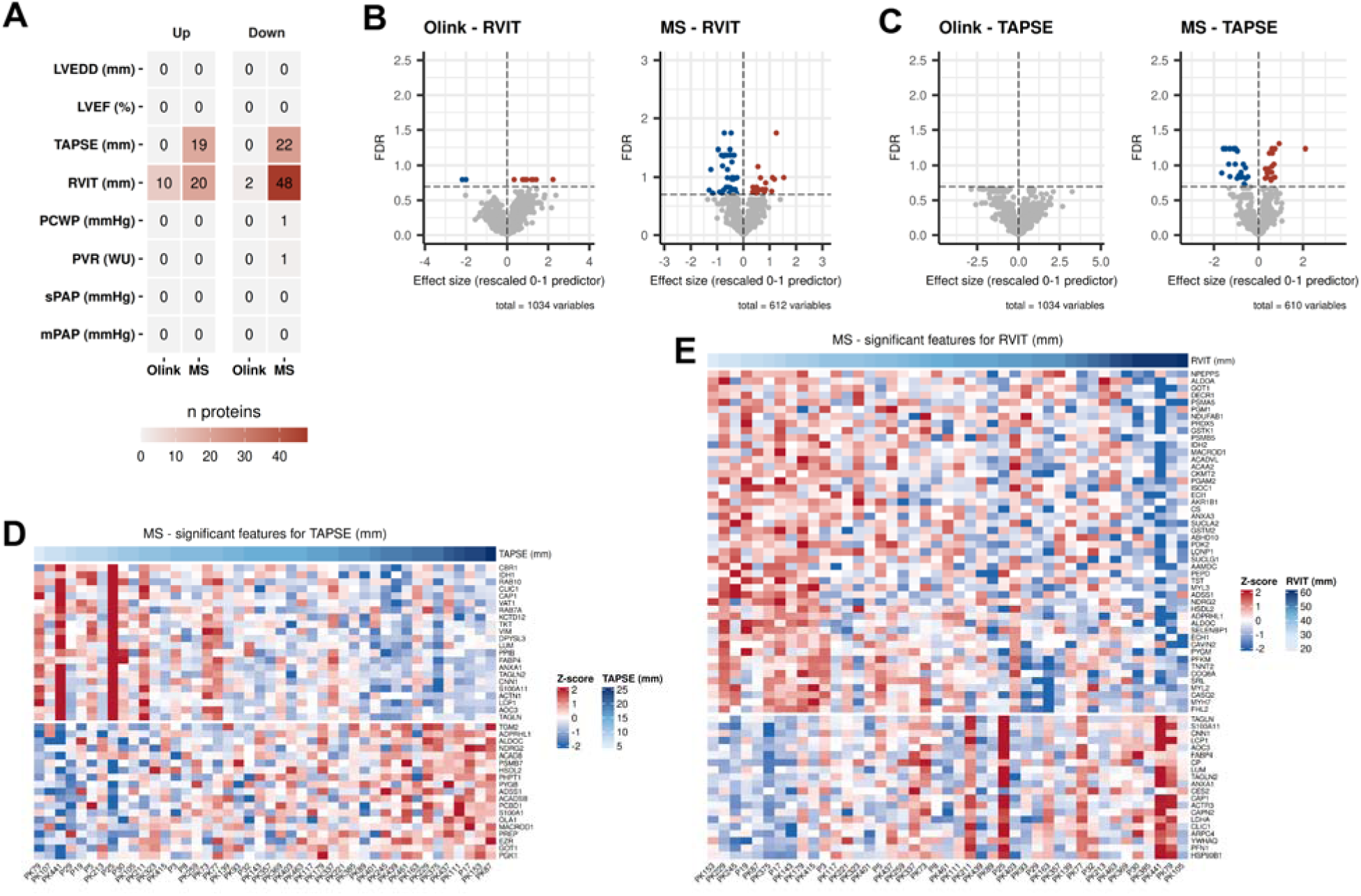
Differential abundance analysis shows that RV tissue proteomics is associated with RV function and size rather than pulmonary pressure load. **A**, Number of proteins with increased or decreased abundance in association with each clinical parameter in the LC MS/MS and Olink datasets. **B-C**, Volcano plots showing protein abundance associations with RVIT (**B**) and TAPSE (**C**) in LC-MS/MS and Olink datasets. Significant proteins were defined using an FDR threshold of 0.2. **D-E**, Heatmap of LC MS/MS proteins significantly associated with TAPSE (D) and RVIT (E). Columns represent individual samples and rows represent proteins.

